# GTPase-powered progressive contraction of a supramolecular ring driving chloroplast division

**DOI:** 10.64898/2026.03.04.709689

**Authors:** Yamato Yoshida, Yuko Mogi, Haruko Kuroiwa, Tsuneyoshi Kuroiwa

**Affiliations:** Department of Biological Sciences, Graduate School of Biological Science, The University of Tokyo; Tokyo 113-0033, Japan; Japan Science and Technology Agency (JST), FOREST; Saitama 332-0012, Japan; Department of Chemical and Biological Science, Faculty of Science, Japan Women’s University; Tokyo 112-8681, Japan

## Abstract

The chloroplast division machinery, known as the division ring, is a supramolecular complex composed of bacterial– and host-derived proteins^1–5^. However, how the division ring generates the force required to sever a chloroplast remains poorly understood. Here, we established an *in vitro* assay in which chloroplasts isolated from *Cyanidioschyzon merolae*^6,7^ undergo GTP hydrolysis–dependent division. Using this assay, we show that Dnm2, rather than FtsZ, acts as the motor driving GTPase-powered contraction of the division ring, thereby physically dividing the chloroplast. We further demonstrate that the division ring is assembled through coiling of a glycosyltransferase-mediated filament and is crosslinked by dimerization of the Dnm2 GTPase domain. Following GTP hydrolysis–dependent force generation, Dnm2 retains its dimeric form in GDP-bound and nucleotide-free states, providing a locking step that suppresses back-slippage of the coiling ring during division. Thus, this mechanical design enables progressive, ratchet-like constriction of the division ring through coiling, overcoming the mechanical load posed by the chloroplast and generating the force required for fission, consistent with quantitative simulations. These findings suggest that a specialized division-ring mechanism, distinct from vesicle fission systems, evolved to mediate endosymbiont fission, allowing host control of endosymbiont proliferation and promoting faithful inheritance of the emerging organelle.

## Introduction

Photosynthesis is one of the few mechanisms by which living organisms can harness energy directly from abiotic sources in the universe^8^. Thus, the acquisition of a photosynthetic organelle grants eukaryotic cells energetic autonomy, liberating them from dependence on external organic carbon sources. The plastid, or chloroplast, is the most widespread photosynthetic organelle and evolved from a free-living cyanobacterial ancestor through an ancient endosymbiotic event^9,10^.

Because of its evolutionary origin via endosymbiosis, the chloroplast cannot form *de novo* in eukaryotic cells and instead proliferates through binary fission^1,4,11,12^. Its division is orchestrated by a supramolecular complex known as the chloroplast division machinery or division ring, which consists of the bacterium-derived protein FtsZ and host-derived components^2,3,11^, including glycosyltransferase PDR1-mediated filaments and a dynamin-related GTPase^13,14^ (Supplementary Fig. 1a). However, the mechanism by which this division ring generates the force required to physically divide the large organelle remains poorly understood.

In bacteria, FtsZ is a key division protein that forms a ring structure, known as the FtsZ ring, at the mid-cell region just prior to cytokinesis^15,16^. Similarly, in chloroplasts, FtsZ assembles into a ring on the stromal side of the inner envelope membrane, marking the initial phase of chloroplast division ring assembly^17–19^. Accordingly, loss of FtsZ function results in defective ring formation and complete failure of chloroplast division^20^. Although the chloroplast FtsZ ring is presumed to generate the constriction force required for division^21^, it is insufficient to do so on its own^22,23^, in contrast to its bacterial counterpart^24,25^.

The plastid-dividing (PD) ring was first characterized as a distinct ring structure involved in chloroplast division in a unicellular alga by electron microscopy^26,27^. Since the initial discovery, PD rings have been identified in various photosynthetic organisms across multiple lineages, including algae and land plants^28–30^. Electron microscopy has revealed several types of PD ring structures with distinct positions relative to the double-envelope membranes, namely the inner, middle, and outer PD rings^31^. Among these, the outer PD ring is notable for its structural robustness. It assembles on the cytosolic side of the outer envelope membrane and consists of a bundle of nanofilaments, each approximately 5–7 nm in diameter, suggesting that it serves as a primary structural component of the chloroplast division-ring complex^13,32^. Proteomic analysis of the division ring revealed that the outer PD ring filament is composed of polyglucan and identified PDR1 as the glycosyltransferase responsible for its synthesis^14^. The glycosyltransferase domain of PDR1 belongs to the GT8 family and shows sequence similarity to glycogenin, which functions as a priming enzyme in glycogen synthesis^33^. Although a homolog of PDR1, MDR1, has been identified in the mitochondrial division machinery of *C. merolae*^34^, bona fide PDR1 orthologs in other photosynthetic eukaryotes remain elusive.

Because of its molecular properties, the PD ring is unlikely to generate the driving force required for chloroplast division. Instead, a chloroplast division–associated dynamin-related protein (cpDRP) has been identified as a GTPase component of the division ring and is thought to contribute to the constriction process^13,22,35^. The term cpDRP refers to dynamin-like GTPases that are thought to be functionally conserved across species and known by different names, such as Dnm2 in *Cyanidioschyzon merolae* and ARC5 (also known as DRP5B) in *Arabidopsis thaliana*.

The dynamin superfamily is involved in various membrane fission events in both eukaryotes and prokaryotes^36–39^. Among its members, classical dynamin is the most extensively studied, particularly for its role in vesicle scission^38–40^. Dynamin can oligomerize into ring-like or helical structures on membranes, and these helical assemblies undergo GTPase-driven constriction^39,41–43^ (Supplementary Fig. 1b). Several models have been proposed to explain this constriction process, with GTPase domain–to–GTPase domain dimerization considered a key mechanistic feature of dynamin as a mechanoenzyme^44–48^.

Structural studies using both full-length and truncated forms of classical dynamin have provided important insights into its mechanical properties^42–44^. However, findings from dynamin studies cannot be directly extrapolated to cpDRPs. One notable difference between cpDRPs and classical dynamin is the extent to which they form filament-like structures at the fission site. Classical dynamin forms continuous, polymerized filaments that encircle the division site^41^, whereas cpDRPs localize as discontinuous patches along the chloroplast division site^22,35^ (Supplementary Fig. 1c). Moreover, the size of the target membrane-enclosed compartment differs markedly between single-membrane vesicles (∼50 nm in diameter) and chloroplasts (∼3,500 nm in diameter in *C. merolae* and ∼7,000 nm in diameter in *A. thaliana*), suggesting that the mechanics of chloroplast fission differ from those of vesicle fission (Supplementary Fig. 1a and b). Although the amino acid lengths of human classical dynamin (864 aa) and cpDRPs (962 aa for Dnm2 and 777 aa for ARC5) are comparable, they share only the ∼300-amino-acid GTPase domain^49^. Classical dynamin can tether to lipid membranes via its pleckstrin homology (PH) domain^38^, whereas cpDRPs are known to associate with outer PD ring filaments and membrane-associated proteins, rather than directly with the chloroplast outer membrane^13,35^. These structural and functional differences, along with the still-unknown mechanism of constriction in the chloroplast division ring, underscore the need for a fundamentally new model of ring constriction.

Because of its cellular simplicity and biological features, the unicellular alga *C. merolae* offers outstanding advantages for studying chloroplast division^6,7^. Each *C. merolae* cell contains only one nucleus, one mitochondrion, and one chloroplast (Supplementary Fig. 2a and b). In addition to synchronization of the cell division cycle, the timing of chloroplast division can also be synchronized by light–dark cycle illumination^50–52^. Owing to this simplified cellular architecture, *C. merolae* remains the only known organism from which dividing chloroplasts with intact division rings can be isolated^28^. Furthermore, we previously established methods to isolate intact division rings from chloroplasts^13^.

In this study, we established an *in vitro* chloroplast division assay using isolated chloroplasts and division rings, enabling detailed mechanistic and kinetic investigations of ring constriction. Combined with imaging, mutagenesis, and nucleotide state-dependent analyses, these studies clearly demonstrated that the GTPase activity of Dnm2 is a fundamental and critical factor in generating the driving force for ring constriction. Finally, we propose that repetitive cycles of Dnm2 GTPase activity, which include GTP binding, hydrolysis, and the resulting conformational changes, together with dimerization and dissociation, promote the sliding and coiling of the polyglucan-based PD ring filaments. This process gradually reduces the ring’s circumference and generates the mechanical force required for chloroplast fission. These findings shed light on a fundamental principle underlying the mechanism of chloroplast division.

## Results

### Kinetics of division-ring constriction during chloroplast division

For a mechanistic understanding of chloroplast division by the chloroplast division machinery (hereafter referred to as the division ring), we first investigated the time-course distribution of its components during chloroplast division in *C. merolae* cells. A gene expression cassette encoding the fluorescent protein Venus fused to Dnm2 under the control of its native promoter, was introduced into *C. merolae* cells via homologous recombination, replacing the endogenous *Dnm2* gene. In transformants expressing Venus-Dnm2, the fusion protein localized as a single ring on the chloroplast during the division phase, and no abnormal phenotype was observed (Fig. 1a).

**Fig. 1.**
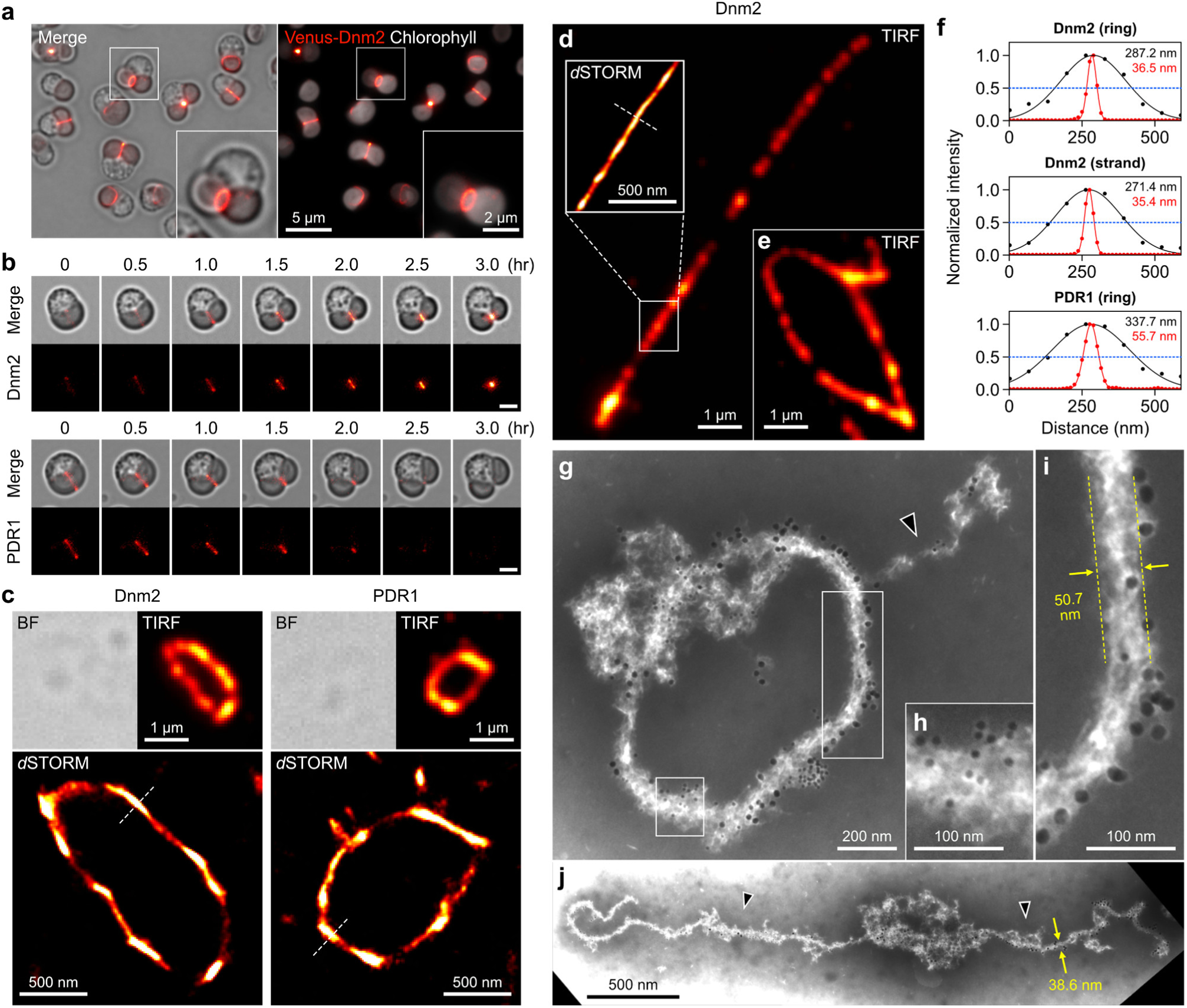
Higher-order architecture and constituent structures of the chloroplast division ring. **a**, Fluorescence images of cells expressing Venus-fused Dnm2. Red, Venus-Dnm2; gray, chlorophyll autofluorescence. **b**, Time-lapse imaging of cells expressing Venus-Dnm2 (upper) and Venus-PDR1 (bottom) during chloroplast division. Scale bars, 2 μm. **c**–**f**, Isolated division rings visualized by TIRF and *d*STORM microscopy. Dnm2 or PDR1 in the isolated rings was detected by immunofluorescence microscopy. The isolated division rings in (**c**) retain a circular structure, whereas those in (**d**) and (**e**) are completely unfolded into a single long filament (**d**) or expanded into an enlarged circular structure (**e**). Cross-sectional intensity profiles of the images in (**c**) and (**d**) along the indicated lines are shown in the graphs (**f**). Gaussian fitting curves and calculated FWHM values are shown in black for TIRF and in red for *d*STORM. **g**–**j**, Electron micrographs of isolated division rings. Dnm2 and PDR1 were immunolabeled with 5-nm and 15-nm gold particles, respectively. Boxed areas in (**g**) are enlarged in (**h**) and (**i**). Arrowheads denote unfolded regions of the isolated division rings.

Time-lapse imaging of individual chloroplasts revealed that the diameter of the division site decreased over time, with a more pronounced reduction during the later stages. Chloroplast division was completed in approximately two to three hours at 42 °C, with constriction rates of 13.2 ± 3.0 nm/min (mean ± SD) in the early phase and 18.7 ± 4.1 nm/min in the late phase, as determined by diameter measurements (*n* = 14) (Supplementary Fig. 2c). The overall constriction rate was 16.5 ± 3.5 nm/min. We next quantified constriction dynamics under a temperature-shift condition in which the incubation temperature was changed from 42 °C to 32 °C during ongoing division. Upon the shift, the constriction rate decreased by approximately half, from 14.6 ± 3.1 nm/min at 42 °C to 7.5 ± 1.8 nm/min at 32 °C (*n* = 5), despite prior assembly of the division ring at the division site (Supplementary Fig. 2c and d). This temperature dependence indicates that division-ring constriction dynamics are governed by enzymatic activity.

As division progressed, the fluorescence intensity of Dnm2 at the midzone of the division ring increased (Fig. 1b, top). Because each Dnm2 molecule was fused to the Venus fluorescent protein, fluorescence intensity served as a proxy for the local copy number of Dnm2 at the ring. These observations suggest that Dnm2 remains associated with the ring with little detectable dissociation and becomes progressively concentrated during constriction.

Notably, the spatiotemporal dynamics of other key components of the division ring, such as PDR1 and FtsZ, differ from those of Dnm2. Our previous study indicated that PDR1 mediates the formation of polyglucan-based PD ring filaments from UDP-glucose and remains associated with these filaments^14^. Consistent with this function, Venus-tagged PDR1 was observed as a single ring on the chloroplast from the onset of division (Fig. 1b, bottom). However, the fluorescence intensity of Venus-PDR1 at the midzone of the division ring remained largely constant during constriction, suggesting that PDR1 does not accumulate as division progresses. Given that its primary role is to synthesize the filament scaffold of the ring, persistent retention of PDR1 in the ring may not be required once constriction begins. FtsZ, a key component on the stromal side of the ring, likewise showed no increase in apparent copy number during constriction (Supplementary Fig. 3a and b). Thus, the spatiotemporal dynamics of Dnm2 during constriction are distinct, supporting an integral role for Dnm2 in driving division-ring constriction.

### Structures of the chloroplast division ring

While the PD ring filament has been considered the primary skeletal element of the division ring^13,14,32^, the persistent localization of Dnm2 during constriction does not exclude the possibility that Dnm2 also contributes to a filamentous scaffold spanning the division ring, in addition to providing the driving force for constriction. To test this possibility and gain insight into the structural organization of the division ring, we compared the protein distributions and ring widths of Dnm2 and PDR1 in isolated division rings using direct stochastic optical reconstruction microscopy (*d*STORM). Division rings were visualized by immunostaining Venus-Dnm2 or Venus-PDR1. We found that total internal reflection fluorescence (TIRF) microscopy yielded consistent images across samples and suggested uniform protein distributions, whereas *d*STORM super-resolution imaging revealed local distortions in the ring and heterogeneous distributions of Dnm2 and PDR1 (Fig. 1c–e).

To quantify these differences, we measured and compared ring widths as full width at half maximum (FWHM) values obtained from Gaussian fits to intensity profiles taken perpendicular to the ring in *d*STORM images. For consistent sampling and to minimize bias from poorly resolved regions, the mean FWHM for each division ring was calculated from ten measurement points with sufficient signal intensity (Fig. 1f). The ring widths were 39.3 ± 3.5 nm for Dnm2 and 51.5 ± 6.0 nm for PDR1, indicating that Dnm2 has a significantly narrower spatial distribution than PDR1 (*p* = 5.51 × 10^−5^) (Fig. 1f, top and bottom). Together, these results suggest that Dnm2 occupies a narrower zone of the division ring than PDR1.

We next examined completely unfolded division rings that were linearly stretched on a glass slide by applying gentle flow through filter paper blotting (Fig. 1d). The mean length of the unfolded rings was 10.8 ± 3.4 μm (*n* = 5), exceeding the average circumference of the *C. merolae* chloroplast (∼9 μm). Notably, the mean width in structurally well-defined linear regions was 36.9 ± 2.8 nm (Fig. 1f, middle), comparable to the width of Dnm2 observed in the circular form (Fig. 1f, top). In addition, the immunofluorescence signal for Dnm2 appeared as discrete puncta aligned along the linearized ring (Fig. 1d). A similar punctate pattern was also observed in partially unfolded samples that retained a circular geometry (Fig. 1e), indicating that this localization pattern is unlikely to be an artifact of ring collapse. Thus, these imaging results support the view that Dnm2 forms oligomeric assemblies and argue against the formation of a continuous filamentous structure within the division ring.

These spatial localizations of Dnm2 and PDR1 were supported by immunoelectron microscopy (immuno-EM) (Fig. 1g–i). While PDR1 was distributed throughout the PD ring bundle (Fig. 1h), Dnm2 was detected on the surface of the outer PD ring bundle (Fig. 1i). This arrangement is consistent with the *d*STORM results showing that Dnm2 occupies a narrower zone than PDR1.

Immuno-EM images of unfolded division rings further indicated that the ring is assembled by coiling. In some isolated division rings, partial unfolding was observed from one or both ends (Fig. 1g,j). Bundle thickness differed markedly between the circular region and the unfolded region, with values of 54.5 ± 9.9 nm (*n* = 10) and 37.2 ± 6.7 nm (*n* = 11; *p* = 2.60 × 10^−4^).

Taken together, these observations suggest that the division ring is built by coiling of a primary filamentous unit composed of PDR1-mediated PD ring filaments with a bundle thickness of ∼37 nm. Dnm2 binds as discrete clusters along the edges of these filaments and localizes to the surface of the coiled structure. This distribution argues against Dnm2 forming a continuous filamentous scaffold within the division ring. On this basis, constriction may proceed through sliding and progressive coiling of a long PD ring unit.

### *in vitro* chloroplast division

The retention of Dnm2 on the surface of the division ring during constriction suggests that its GTPase activity provides the primary driving force for constriction. To assess whether elevating Dnm2 GTPase activity promotes chloroplast division, we incubated isolated chloroplasts bearing division rings with 2.5 mM GTP at 42 °C to enhance Dnm2 GTPase activity. Time-lapse imaging showed that the constriction site gradually narrowed over 12 hours and culminated in chloroplast division (Fig. 2a and Supplementary Video S1). By contrast, no constriction was observed after 12 hours in the absence of GTP, indicating that *in vitr*o chloroplast division is driven by a GTP-dependent reaction (Fig. 2b and Supplementary Video S2). The average GTP-dependent constriction rate, determined from diameter measurements, was 1.66 ± 0.35 nm/min (*n* = 5 individual chloroplasts) (Fig. 2c). Imaging from an orthogonal viewing angle further corroborated progressive constriction of the division ring (Fig. 2d). Furthermore, GTP-stimulated constriction was also observed in isolated division rings (Fig. 2e and Supplementary Video S3). As constriction proceeded, the fluorescence intensity of Venus-Dnm2 increased in the mid-zone, consistent with *in cellulo* observations (Fig. 1b, top). However, the constriction rate was significantly lower. It was approximately one-tenth of the rate observed *in cellulo* (Supplementary Fig. 2c).

**Fig. 2.**
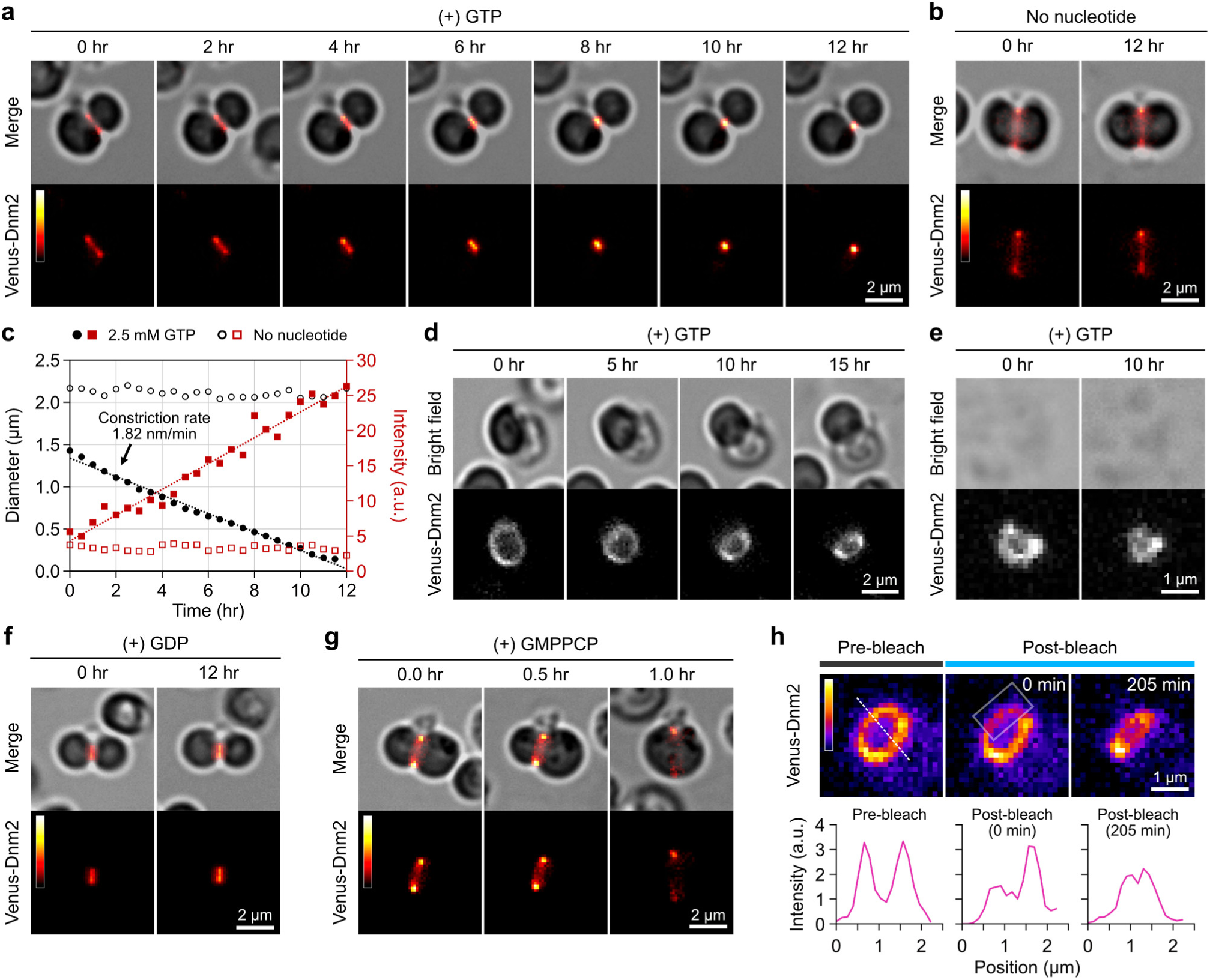
GTPase-driven contraction of the division ring leading to fission of isolated chloroplasts. **a**,**b**, Time-lapse imaging of the division of isolated chloroplasts in front view in the presence (**a**) and absence (**b**) of 2.5 mM GTP. See also Supplementary Videos S1 and S2. **c**, Measurements of chloroplast division-plane diameter (black) and Venus–Dnm2 fluorescence intensity at the mid-region (red) for the same chloroplast as shown in (**a**), in the presence and absence of 2.5 mM GTP. **d**, *In vitro* chloroplast division in side view. **e**, *In vitro* contraction of an isolated membrane-free division ring in the presence of 2.5 mM GTP. See also Supplementary Video S3. **f**,**g**, Effects of GTP analogs on the division ring in isolated chloroplasts. See also Supplementary Video S4. **h**, Photobleaching analysis of the constricting division ring on an isolated chloroplast in the presence of 2.5 mM GTP. Cross-sectional intensity profiles along the indicated lines are shown in the graphs. The boxed region indicates the photobleached area.

Because the *in vitro* experiments were performed with isolated intact chloroplasts, GTPases associated with the division machinery, including FtsZ2A and FtsZ2B in addition to Dnm2, should remain within the chloroplast. However, although Venus-tagged FtsZ2B forms a ring at the division site *in cellulo* (Supplementary Fig. 3a,b), no FtsZ ring was detected in isolated chloroplasts (Supplementary Fig. 3c,d). Instead, the FtsZ2B-Venus signal was dispersed throughout the stroma, suggesting that exogenously added GTP might not effectively access the stromal compartment. These observations indicate that the FtsZ ring may contribute less to constriction during *in vitro* chloroplast division, which could partly explain the reduced constriction rate.

Although the nucleotide-free (apo) condition did not produce a detectable phenotype (Fig. 2b), we next examined whether inhibiting specific steps in the Dnm2 GTP hydrolysis cycle affects *in vitro* chloroplast division. To bias Dnm2 toward distinct nucleotide-bound states, we used GDP to favor a post-hydrolysis-like inactive state and GMPPCP, a non-hydrolyzable GTP analog, to favor a pre-hydrolysis active state. Each nucleotide analog was added to isolated dividing chloroplasts and incubated under standard conditions. GDP did not induce obvious changes in division-ring morphology over 12 hours (Fig. 2f). By contrast, after a 1-hour incubation with GMPPCP, the constricting division ring rapidly relaxed and the constriction site was lost within minutes (Fig. 2g and Supplementary Video S4). Consistent with this rapid loss of the constricting ring, Venus-Dnm2 fluorescence on the ring decreased after incubation. These results indicate that shifting Dnm2 toward a pre-hydrolysis state induces rapid relaxation and uncoiling of the constricting ring. Together, they demonstrate that chloroplast division is driven by physical constriction of the chloroplast envelope against mechanical resistance, including membrane tension.

Thus, these findings indicate that both structural retention and coiling of the division ring are tightly coupled to Dnm2 GTPase activity, and they highlight the need for sufficient mechanical integrity and force generation to sustain progressive constriction during chloroplast division. This requirement for sustained mechanical force may distinguish the chloroplast division ring from fission machineries that drive vesicle scission and other fission events involving small membrane-bound organelles.

As the final part of our *in vitro* chloroplast division analysis, we examined division-ring coiling using a fluorescence recovery after photobleaching (FRAP)-based tracking assay. After photobleaching the Venus-Dnm2 signal, dividing chloroplasts were incubated under GTP-containing conditions to induce constriction (Fig. 2h). Line-profile comparisons across the ring circumference before and after photobleaching showed that the bleached region progressively spread along the ring during constriction. These results indicate that Dnm2 molecules, tethered along the edge of a long filamentous unit, move along the division ring in coordination with constriction. Based on a series of microscopic analyses of unfolded division rings, together with the FRAP-based tracking assay, we propose that this movement is driven by progressive coiling of the long filamentous unit, which is powered by Dnm2 GTPase activity.

### Molecular orientation of Dnm2 on the division ring

Our working model proposes that GTPase activity drives contractile movement of the division ring through structural rearrangements that involve coiling. Realization of the ring’s contractile function requires that Dnm2 molecules be spatially organized in an ordered, directional manner to convert chemical energy into unidirectional motion and constrictive force.

We therefore examined the molecular orientation of Dnm2 on the division ring using polarized fluorescence microscopy. Venus was directly fused to the C-terminal α-helix of Dnm2 to constrain fluorophore orientation, generating Dnm2-conVenus (Supplementary Fig. 4a). Cells expressing Dnm2-conVenus were then imaged by polarized fluorescence microscopy (Supplementary Fig. 4b).

In our setup, the polarizers were aligned along the *X*-axis (horizontal) and *Y*-axis (vertical) in the plane of the microscope stage. Polarized fluorescence imaging of Dnm2-conVenus revealed that the apparent width of the division ring depended on the ring’s orientation. Specifically, when the chloroplast division plane was nearly parallel to the *Y*-axis, the ring appeared consistently narrower through the vertical polarizer than through the horizontal polarizer (Fig. 3a). This difference was confirmed by measuring the FWHM of fluorescence intensity line profiles, yielding values of ∼267 nm (vertical) and ∼367 nm (horizontal) (Supplementary Fig. 4c,d). Conversely, when the division plane was nearly parallel to the *X*-axis, the relationship was reversed (Fig. 3b). The apparent ring width varied periodically with the rotation angle of the division plane, following a sine dependence consistent with a projection effect (Fig. 3c). This polarization-dependent behavior persisted in isolated chloroplasts and in cells during the late stage of chloroplast division (Supplementary Fig. 4e,f). By contrast, when the division ring was imaged from an orthogonal (side) view, the polarized fluorescence pattern showed no detectable change (Supplementary Fig. 4g).

**Fig. 3.**
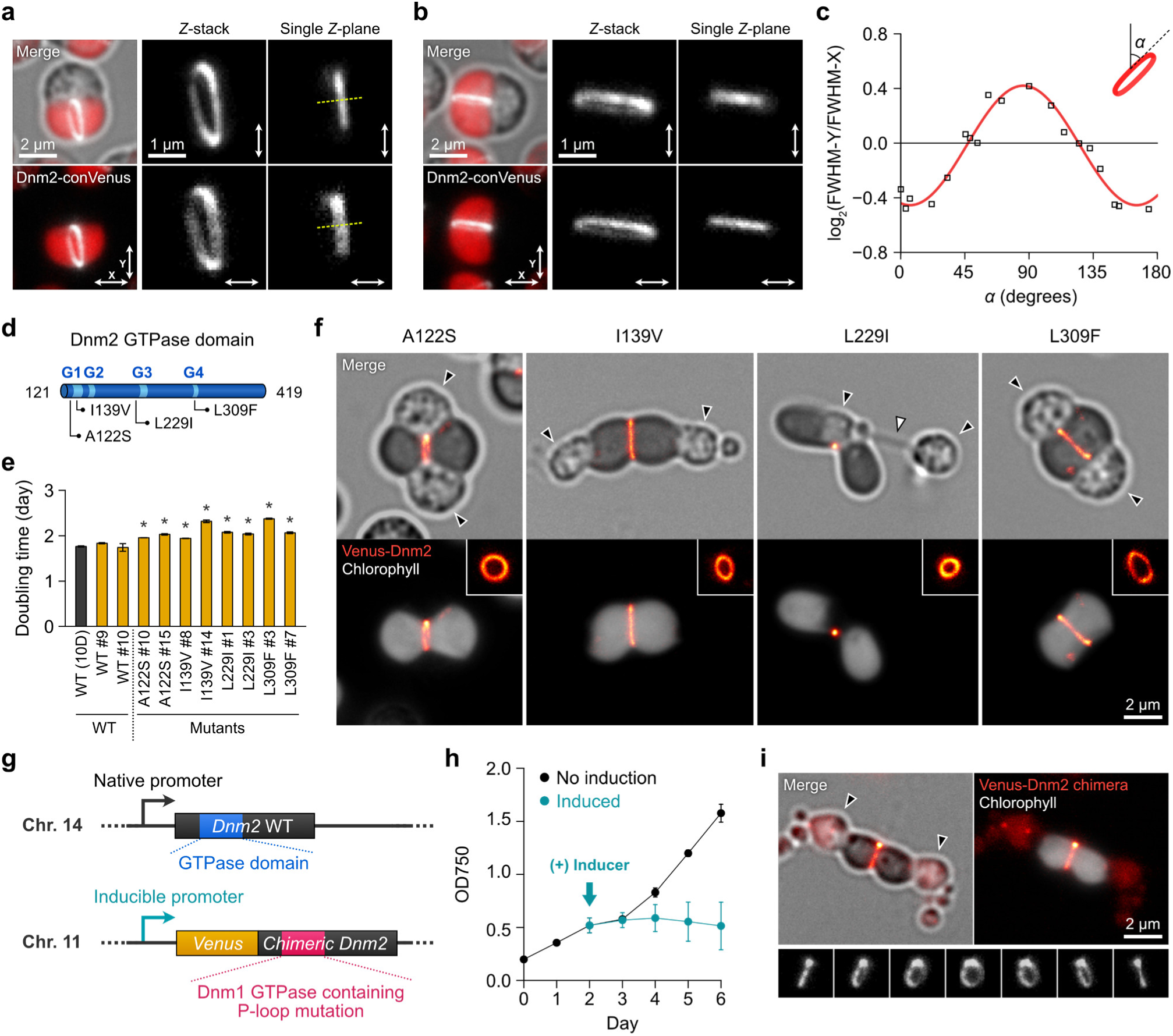
Ordered Dnm2 localization and GTPase-dependent constriction of the chloroplast division ring. **a**,**b**, Polarized fluorescence microscopy of Venus-tagged Dnm2 with a constrained molecular orientation (Dnm2-conVenus). Images were acquired using parallel polarizers aligned along the *X*– and *Y*-axes (indicated by bidirectional arrows). Cross-sectional intensity profiles of the images in (**a**) along the indicated lines are shown in Supplementary Fig. 4. **c**, Log₂ ratio of FWHM (*Y*/*X*) plotted as a function of rotation angle (α). The red curve indicates the best-fit sine function. **d**, Schematic of the G1–G4 motifs within the Dnm2 GTPase domain (residues 121–419) and the positions of the amino acid substitutions. **e**, Doubling times of wild-type (WT) and mutant transformants. Yellow bars indicate cell lines expressing Venus-tagged Dnm2. Data shown represent mean ± standard error; \**P* < 0.05 for wild-type versus mutant Venus-Dnm2, as determined by *t*-test. **f**, Representative fluorescence images of transformants carrying the indicated mutations. Insets show side views of the division rings. Black arrowheads indicate the putative nuclear position, and the white arrowhead marks the spindle. **g**, Schematic of the endogenous *Dnm2* locus and expression constructs for native Dnm2 and an inducible, promoter-driven Venus-tagged chimeric Dnm2 (Venus-Dnm2 chimera) harboring a Dnm1-derived GTPase domain with a P-loop mutation (K→A). **h**, Growth curves of cells expressing the Venus-Dnm2 chimera under non-induced (black) and induced (green) conditions. The arrow indicates the time point at which the culture medium was replaced with nitrate medium to induce expression. **i**, Representative fluorescence image of a cell expressing the Venus-Dnm2 chimera.

Thus, these polarization-dependent width changes suggest that Dnm2 molecules in the mid-zone of the division ring are oriented approximately perpendicular to those in the outer zones. This in turn indicates that a defined molecular orientation is confined to Dnm2 molecules positioned at the interface with the coiling filament.

### Mutagenesis of Dnm2’s GTPase domain

To assess the functional importance of the Dnm2 GTPase domain in division-ring constriction dynamics, we generated a Venus-Dnm2 variant carrying a single amino acid substitution within the GTPase domain. Mutations in the well-characterized G1–G4 motifs were lethal, and we were unable to obtain stable transformant lines. Therefore, to preserve intrinsic GTPase activity and minimize structural changes, we introduced a conservative substitution within or adjacent to the G1–G4 motifs (Fig. 3d). The substituted position was chosen based on a multiple sequence alignment of 100 Dnm2 homologs from diverse organisms, in which the alternative residue is conserved in a subset of homologs (Supplementary Table 1).

We found that the A122S, I139V, L229I, and L309F substitutions prolonged the doubling time (Fig. 3e). Microscopic observation revealed that the delayed proliferation in these mutants was associated with defects in chloroplast division, although the onset varied depending on the substitution site. While the division ring assembled normally at the division site, constriction was arrested or delayed in mutants carrying A122S, I139V, or L309F (Fig. 3f). As a representative example, the constriction rate in I139V mutant cells was 10.4 ± 2.2 nm/min (*n* = 11), significantly lower than in wild type (16.5 ± 3.5 nm/min, *p* = 2.2×10^−3^) (Supplementary Fig. 2c, right). In these mutants, more than half of cells underwent both chloroplast and cell division normally, whereas the remaining cells exhibited failed or aberrant chloroplast division accompanied by cytokinesis failure. The incomplete coordination between chloroplast division and cell division, likely reflecting stochastic failures in chloroplast division, may underlie the prolonged doubling time in these mutants. In contrast, although two apparently separate chloroplasts were observed in dividing L229I mutant cells, the persistent localization of a constricted division ring at the mid-region of the daughter chloroplasts suggested incomplete membrane scission and continued connectivity between the two chloroplast compartments (Fig. 3f). Consequently, chloroplast inheritance was unequal, and after nuclear segregation one daughter cell often lacked a chloroplast.

These results highlight the importance of the Dnm2 GTPase domain and GTPase activity in the mechanism underlying division-ring constriction. To further dissect Dnm2 function within the division ring and identify where force is applied, we constructed a chimeric Dnm2 in which the GTPase domain was replaced with that of Dnm1, a DRP required for mitochondrial division in *C. merolae*^53^ (Fig. 3g and Supplementary Fig. 5). In addition, we introduced a lysine-to-alanine substitution in the P-loop motif to impair GTPase activity, analogous to the well-characterized K44A mutation in classical dynamin. The Venus-tagged chimeric Dnm2 was co-expressed with endogenous Dnm2 in transformant cells under the control of an inducible promoter. This system allowed us to assess whether the chimera, bearing a Dnm1-derived GTPase domain, localizes to the chloroplast division ring and whether it interferes with endogenous Dnm2-dependent ring constriction.

As a result, transformant cells grew normally under non-inducing conditions, whereas induction of the chimeric Dnm2 impaired cell growth and caused defects in chloroplast division (Fig. 3h,i), similar to those observed in the single–amino acid substitution mutants (Fig. 3f). We also confirmed that the chimeric Dnm2 localized to the chloroplast division site and formed a ring similar to that formed by wild-type Venus-Dnm2. These findings indicate that replacing the Dnm2 GTPase domain with the Dnm1 GTPase domain and disrupting GTPase activity do not compromise targeting to the division ring, suggesting that regions outside the GTPase domain mediate localization to the division ring. Moreover, incorporation of a catalytically inactive GTPase domain into the ring severely disrupted constriction, despite the presence of endogenous Dnm2 in the transformed cells.

These results suggest that division-ring targeting is mediated by regions outside the GTPase domain and is independent of both the origin and catalytic activity of the GTPase domain. In contrast, the strong dominant-negative effect of a catalytically inactive GTPase domain indicates that cooperative action among homologous GTPase domains within the ring is required to generate the driving force for constriction.

### Nucleotide-state–dependent GTPase-domain dimerization

A series of site-directed mutagenesis and domain-replacement experiments suggests that the Dnm2 GTPase domain generates the driving force for ring constriction. Classical dynamins and several DRPs form homodimers via such GTPase-domain interactions during GTP hydrolysis, suggesting that Dnm2 likely adopts a similar dimeric configuration. To directly test this possibility and evaluate the intrinsic GTPase activity of Dnm2, we constructed truncated proteins containing the GTPase domain and its adjacent helices (designated GD-2H). Because an additional C-terminal helix contributes to formation of the bundle signaling element (BSE) in classical dynamins^44,54^, we also designed an extended construct, GD-3H, in which this helix is appended via a 15-amino-acid flexible linker to mimic the BSE (Fig. 4a,b).

**Fig. 4.**
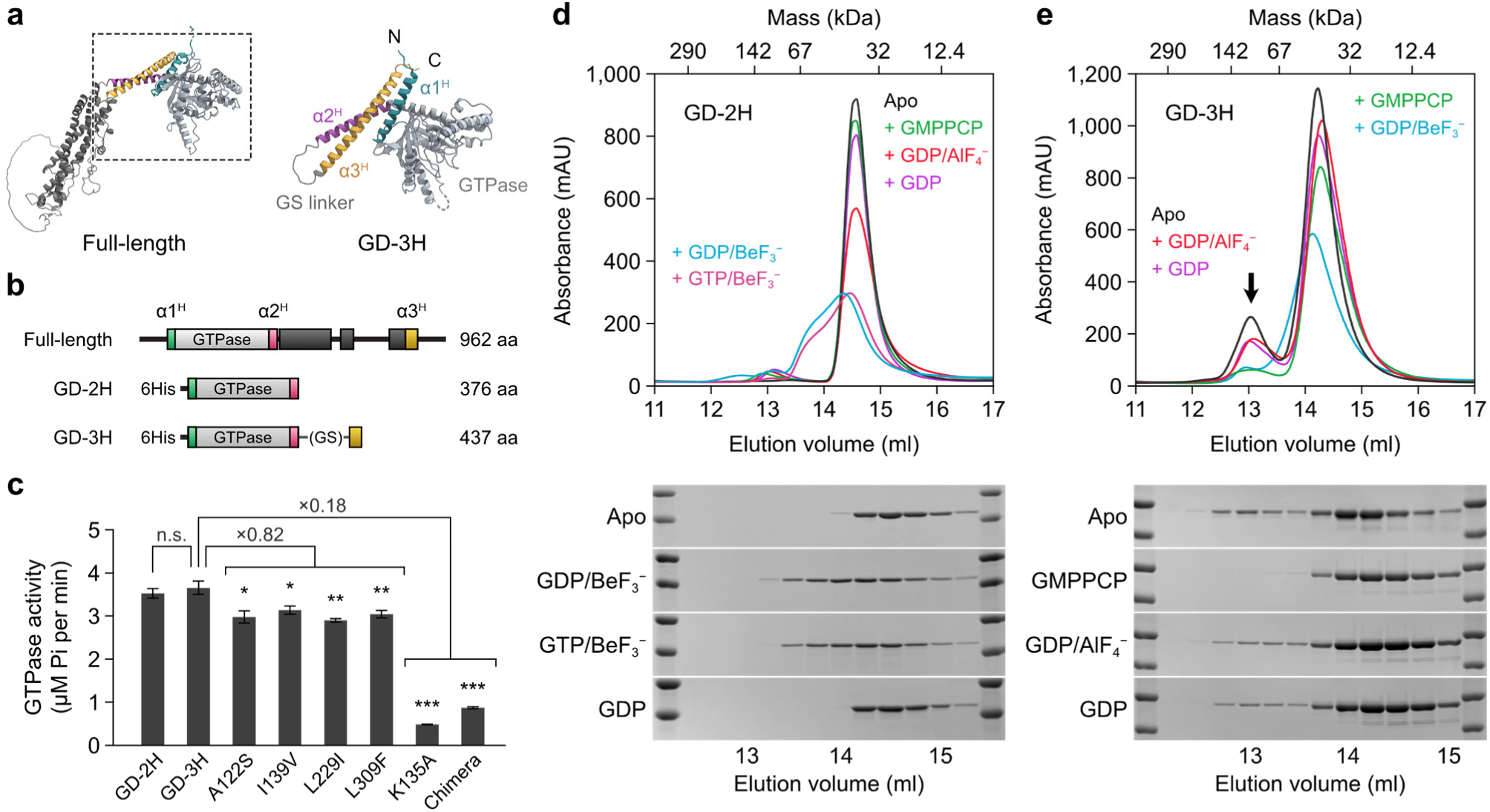
Nucleotide state–dependent GTPase dimerization of Dnm2. **a**, Simulated protein structures of full-length Dnm2 and its GTPase domain (GD-3H). **b**, Schematic representations of domain compositions of full-length Dnm2, GD-2H, and GD-3H. Putative three α-helices transmitting conformational changes derived from GTP hydrolysis are shown in green (α1^H^), magenta (α2^H^), and yellow (α3^H^). **c**, GTPase activities of the Dnm2 GTPase domains. 0.5 μM of protein was used for each sample. **d**,**e**, Size-exclusion chromatography of the Dnm2 GTPase domain in the presence of different guanine nucleotide analogs. Dnm2 GTPase (1 mg/mL) was incubated with each analog for 60 min at 42 °C and then injected onto an ENrich SEC650 column. Retention volumes for the monomeric and dimeric forms of apo-state GD-3H were 14.3 mL and 13.0 mL, respectively. Retention volumes of molecular-weight standards are shown above. Left and right lanes in each SDS-PAGE analysis correspond to molecular-weight standards (upper band, 50 kDa; lower band, 37 kDa).

The GTPase activities of GD-2H and GD-3H were approximately 3.52 min⁻¹ and 3.65 min⁻¹, respectively, expressed as μmol Pi released per minute per μmol of protein, and no statistically significant difference was observed between the two constructs (Fig. 4c). These activities are substantially higher than the activity reported for a truncated form of human classical dynamin under a comparable condition^44^. To further examine the contribution of conserved residues to GTPase activity, we tested GD-3H constructs carrying point mutations. As expected, both the K135A mutant and the chimeric Dnm2 containing a P-loop mutation showed a marked reduction in GTPase activity, to 0.13-fold and 0.24-fold of the wild-type GD-3H level, respectively. By contrast, constructs harboring conservative substitutions (A122S, I139V, L229I, and L309F) exhibited statistically significant but modest decreases, ranging from 0.79– to 0.86-fold relative to the wild-type GD-3H.

Using the truncated wild-type GD-2H and GD-3H constructs, we next performed size-exclusion chromatography (SEC) to assess their oligomeric states with and without nucleotides. GD-2H remained monomeric in the apo state as well as in the presence of GMPPCP or GDP (Fig. 4d). Notably, under conditions that mimic the transition state of GTP hydrolysis (GDP/BeF₃⁻ or GTP/BeF₃⁻), the monomer peak was partially diminished and broadened toward higher apparent molecular weights, although it did not reach the expected molecular weight of a dimer. These results indicate that the peak broadening likely reflects subtle conformational changes and/or transient self-association.

We also examined GD-3H and observed behavior distinct from that of GD-2H (Fig. 4e). Notably, GD-3H displayed an apparent dimer peak in the apo state in addition to the monomer peak. This dimer peak was also observed in the presence of GDP and under a GDP·Pi transition-state mimic generated with GDP/AlF₄⁻. By contrast, the dimer peak was reduced in the presence of GMPPCP or under a GTP-like pre-hydrolysis state mimic generated with GDP/BeF₃⁻.

Because only GD-3H exhibited an apparent dimer peak, the C-terminal helix likely promotes GTPase-domain dimerization, consistent with previous findings for canonical dynamins. However, the ability of GD-3H to form dimers across multiple stages of the GTP hydrolysis cycle, including the apo state, appears to differ from that of canonical dynamins and other DRPs. Moreover, the reduced dimer population of GD-3H in the presence of GMPPCP (Fig. 4e) may provide a molecular explanation for the GMPPCP-induced relaxation of the constricting ring in isolated chloroplasts (Fig. 2g).

### Mechanistic model of force generation in the chloroplast division ring

This property of Dnm2 may reflect the specific mechanical requirements of chloroplast division, as chloroplasts have a much larger volume and greater mechanical resistance than vesicles. Results from our *in vitro* chloroplast division assays (Fig. 2) and SEC analyses of GD-3H (Fig. 4e) suggest that sustained ring constriction requires a large fraction of Dnm2 molecules to repeatedly hydrolyze GTP and form GTPase-domain dimers, thereby maintaining tension and counteracting mechanical load. Accordingly, the ability to dimerize across multiple stages of the GTP hydrolysis cycle could facilitate efficient constriction of the chloroplast division ring.

Based on these results, we propose a model in which Dnm2 acts as the principal actuator driving stepwise, contractile constriction of the division ring (Fig. 5a). In this model, Dnm2 molecules do not assemble into a continuous filament (Fig. 1d, Supplementary Fig. 1c). Instead, Dnm2 localizes to the edges of PDR1-mediated filament units with a constrained orientation and associates with these units via its non-GTPase regions (Fig. 1i; Fig 3a–c; Fig 3i). We propose that GTPase–GTPase dimerization occurs between opposing Dnm2 molecules on adjacent filament units and may be accompanied by an ∼90° molecular rotation (Fig. 5b). Thus, Dnm2 could establish force-transmission points both at its interfaces with the PDR1-mediated filaments and at sites of GTPase–GTPase dimerization.

**Fig. 5.**
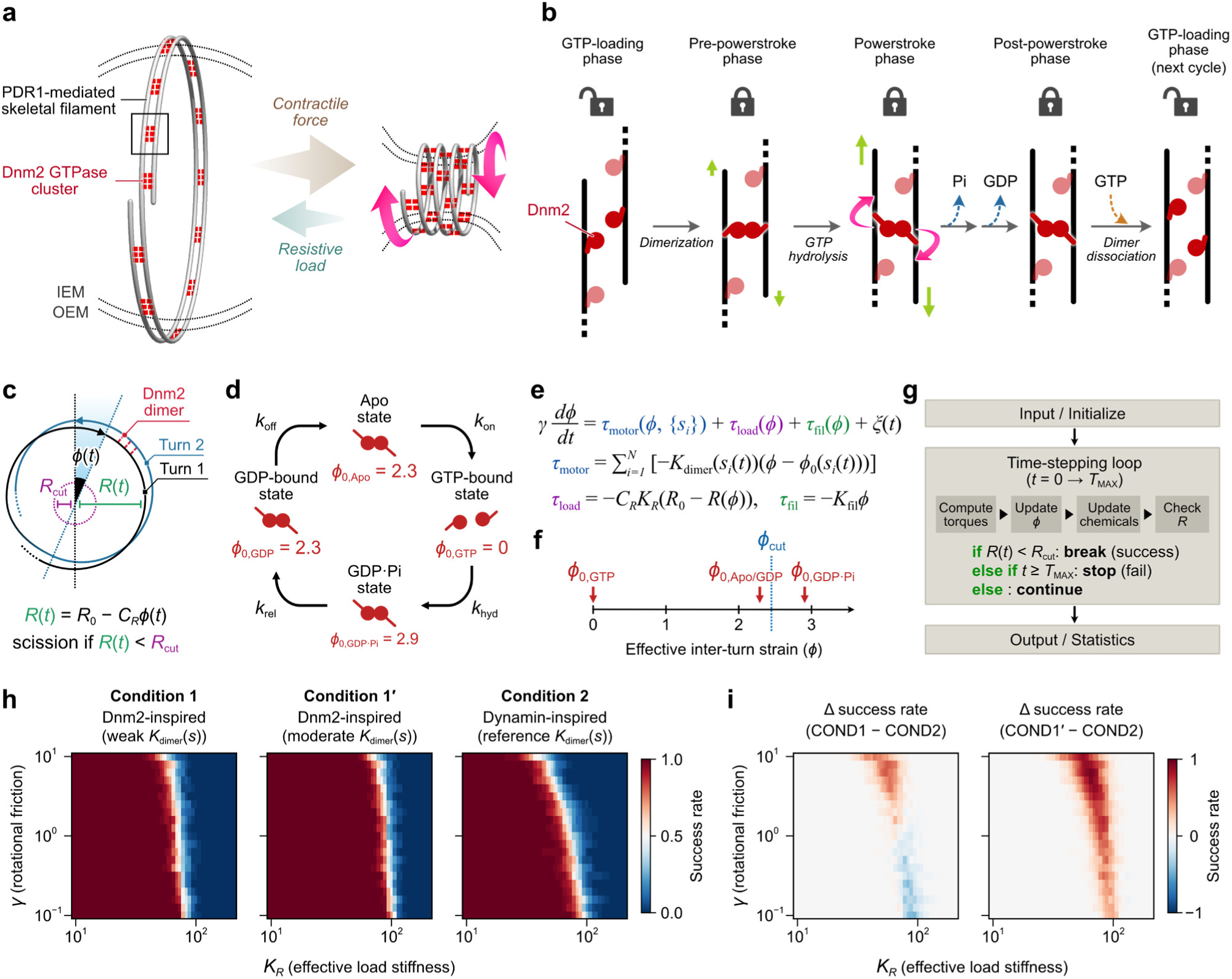
GTPase-powered ring contraction driven by filament sliding and coiling. **a**, Model for GTP hydrolysis–driven ring contraction. Dnm2 clusters bridge PDR1 scaffold filaments, maintaining a coiled division-ring architecture and generating contractile force against the resistive load. **b**, Proposed Dnm2 mechanochemical cycle. GTPase-domain dimerization and a ratchet-like conformational change upon hydrolysis drive unidirectional filament sliding. In these schematics, only Dnm2 molecules localized within the interfilament space are shown. **c**, Minimal coarse-grained model for GTPase-driven, coiling-mediated constriction of the division ring. The collective action of many motors is coarse-grained into a single effective twist coordinate, *ϕ*(*t*), which maps linearly onto the neck radius as *R*(*t*) = *R*_0_ − *C_R_ϕ*(*t*). **d**, State-dependent motor engagement in the coarse-grained model. Motor activity is parameterized by *ϕ*_0_(*s*) (preferred twist) and *K*_dimer_(*s*) (coupling stiffness) across the chemical states. See also Supplementary Fig. 6. **e**, Overdamped Langevin equation for the coarse-grained model. **f**, One-dimensional plot of *ϕ* for each chemical state relative to the scission threshold *ϕ*_cut_. **g**, Workflow of the coarse-grained mechanochemical simulations. **h**, Scission success-rate phase diagrams for the Dnm2-inspired schemes (Conditions 1 and 1′) and the dynamin-inspired scheme (Condition 2). **i**, Differential success-rate map (Δ success rate) summarizing the parameter regime in which post-hydrolysis coupling improves scission robustness.

As shown by a series of SEC analyses, Dnm2 dimers dissociate into monomers upon GTP binding (Fig. 4e). These monomers undergo conformational changes at the onset of GTP hydrolysis (Fig. 4d) and subsequently reassemble into dimers, potentially with new pairing combinations. The dimers remain stable in the GDP-bound and apo states.

Together with our data, the GTPase-domain dimerization and GTP-hydrolysis–driven conformational changes observed in many dynamin-family proteins^46^ suggest that Dnm2 generates force through a similar power-stroke mechanism. In addition, Dnm2’s ability to form dimers not only in a pre-hydrolysis state but also in post-hydrolysis states (GDP-bound and apo) may enable a robust, ratchet-like mechanism that promotes progressive ring constriction.

In turn, repeated GTP hydrolysis by Dnm2 could drive unidirectional sliding and coiling of PDR1-mediated filaments, thereby producing the mechanical work that ultimately divides the chloroplast into two daughter organelles (Fig. 5b).

### Coarse-grained mechanochemical simulation of division-ring constriction

This mechanistic model implies that the dominant torque-generating step occurs in the GDP·Pi state, whereas the GDP– and apo-state phases adopt a torsionally locked configuration that limits strain relaxation, thereby preserving stored torsional strain after nucleotide turnover. To test this idea, we built a minimal model of GTPase-driven, coiling-mediated constriction of the division ring (Fig. 5c, Supplementary Notes). Although the division machinery is geometrically complex, the key readout relevant to chloroplast division is the progressive reduction of the neck radius at the division site. We therefore coarse-grained the collective action of many GTPase-dependent motors into a single effective twist coordinate, *ϕ*(*t*), which maps linearly onto the radius as *R*(*t*) = *R*_0_ − *C_R_ϕ*(*t*). State-dependent motor engagement is captured by *ϕ*_0_(*s*) (preferred twist) and *K*_dimer_(*s*) (coupling stiffness), whereas the organelle membrane and associated constraints are represented as a resistive load with stiffness *K_R_*, subject to rotational friction *γ* (Fig. 5d,e). Accordingly, scission is achieved when *R*(*t*) falls below a critical radius *R*_cut_, equivalently when *ϕ*(*t*) exceeds *ϕ*_cut_ = (*R*_0_ − *R*_cut_)/ *C_R_* (Fig. 5f). In addition, motivated by the partial dimerization of GD-3H observed by SEC, we set *K*_dimer_ in the GDP·Pi/GDP/apo states to one-third (Condition 1) or one-half (Condition 1′) of the GDP·Pi-state value used in a classical dynamin-inspired scheme (Condition 2) (Supplementary Fig. 6a).

With this model and parameterization (Supplementary Table 2), we simulated the dynamics of *ϕ*(*t*) over a range of load stiffness *K_R_* and rotational friction *γ* (Fig. 5g). Scission was scored when *R*(*t*) < *R*_cut_ (equivalently, *ϕ*(*t*) > *ϕ*_cut_). This *K_R_*−*γ* parameter sweep spans division scenarios ranging from single-membrane-bounded vesicles (low *K_R_*, low *γ*; dynamin-based rings) to double-membrane-bounded organelles (high *K_R_*, high *γ*; the chloroplast division ring). The resulting phase diagrams revealed a sharp boundary between scission and non-scission regimes that shifted with increasing *K_R_* (Fig. 5h). Notably, the boundaries diverged at higher *γ*. In the Dnm2-inspired schemes (Conditions 1 and 1′), the scission boundary was largely insensitive to *γ*, whereas in the classical dynamin-inspired scheme (Condition 2), increasing *γ* reduced load tolerance and expanded the non-scission regime.

The qualitative difference between Conditions 1 and 2 was more clearly visualized in phase diagrams of the difference in scission success rate (Δ success rate = Condition 1 − Condition 2) (Fig. 5i, left). At lower *γ* (approximately 0.1–1), the success rate near the boundary is higher in Condition 2 than in Condition 1, yielding a negative Δ success rate. This sign reverses at higher *γ* (approximately 1–10), and Δ success rate becomes positive. The superior performance of the Dnm2-inspired scheme at higher *γ* is consistent with a scenario in which the structural complexity of the chloroplast division ring and associated membrane proteins increases the effective rotational friction. By contrast, the superior performance of the classical dynamin-inspired scheme at lower *γ* is consistent with a low-friction regime in which strong coupling confined to the GDP·Pi state is sufficient to drive scission. Overall, these trends support the model prediction that higher load stiffness (*K_R_*) and higher rotational friction (*γ*) penalize schemes lacking post-hydrolysis coupling, whereas at lower *K_R_* or lower *γ*, strong coupling confined to the GDP·Pi state is sufficient for scission. In addition, a moderate *K*_dimer_ estimate (Condition 1′) improves the performance of the Dnm2-inspired scheme even at lower *γ*, expanding the parameter region in which it outperforms Condition 2 (Fig. 5i, right). Differences between the Dnm2-inspired and classical dynamin-inspired schemes were also evident in the time to scission. Under high-*γ* conditions, the Dnm2-inspired schemes (Conditions 1 and 1′) achieved scission more rapidly (Supplementary Fig. 6b,c).

Taken together, these findings support the model prediction that post-hydrolysis coupling improves robustness under increased load stiffness and rotational friction. They further suggest that chloroplast division requires a highly reliable, coiling-driven contractile mechanism to overcome the structural resistance imposed by chloroplast architecture. A stiff filament scaffold, combined with nucleotide-state–dependent dimerization across multiple phases, provides a built-in locking step that suppresses back-slippage. This design yields a ratchet-like mechanism that drives robust, progressive ring constriction.

## Discussion

### Design principles of the chloroplast division ring

Our results indicate that the core structural framework of the chloroplast division ring comprises PDR1-mediated filament units and the GTPase Dnm2. Dnm2 forms discrete clusters that localize along these filament units. The punctate distribution resembles that observed at the chloroplast division site in *Arabidopsis thaliana*, suggesting that cpDRPs, including Dnm2 and its homologs, do not assemble into continuous filaments, unlike classical dynamin (Supplementary Fig. 1).

As demonstrated by an *in vitro* chloroplast division assay, chloroplasts undergo physical division driven by GTP-dependent contraction of the division ring (Fig. 2a). Mutagenesis and domain-exchange analyses indicate that the contractile force is generated primarily by coordinated Dnm2 GTPase activity (Fig. 3f,i). Furthermore, images of unfolded division rings and micrometer-scale molecular movements tracked by FRAP during constriction support a coiling-driven mode of ring contraction powered by the GTP-dependent activity of Dnm2 (Fig. 1d and Fig. 2h). Taken together, the observations suggest a division-ring design in which the PDR1-mediated filament provides the primary structural scaffold, whereas the Dnm2 GTPase supplies localized, nucleotide-state–dependent force-generating modules (Fig. 5a,b).

The combination of a stiff PDR1-mediated filament scaffold and Dnm2, which undergoes nucleotide-state–dependent dimerization across pre– and post-hydrolysis states (Fig. 4e), constitutes a chloroplast-specific division-ring contractile mechanism distinct from other membrane fission systems. Given the structural complexity of the chloroplast, the effective load and rotational friction during chloroplast division are expected to be substantially higher than during vesicle scission.

Coarse-grained simulations of coiling-driven constriction of the division ring provide a quantitative rationale for the advantage of post-hydrolysis dimerization of Dnm2 in increasing the scission success rate across a broad range of load stiffness and rotational friction (Fig. 5c–i). Together, the data support a power-stroke mechanism in the division ring coupled to a post-hydrolysis locking step that suppresses back-slippage and enables ratchet-like, progressive, coiling-driven ring constriction.

### Functional differences between FtsZ and cpDRP in chloroplast division

The principle of chloroplast proliferation via binary fission is evolutionarily conserved among photosynthetic eukaryotes. However, some factors responsible for chloroplast division are conserved only within a restricted taxonomic range^23,55,56^. Such a phylogenetic distribution suggests an evolutionary trend toward functional refinement of the chloroplast division machinery, promoting more efficient and tightly regulated organelle proliferation. Consistent with this view, several land plant–specific chloroplast division factors have been identified, some of which coordinate chloroplast division frequency with developmental programs^57^.

Among known genes related to chloroplast division, FtsZ and cpDRPs are widely conserved across photosynthetic eukaryotes. Although both proteins bind and hydrolyze GTP, their subcellular localizations and modes of action differ fundamentally. FtsZ acts in the stromal compartment, whereas cpDRPs localize to the cytosolic face of the outer chloroplast envelope. Moreover, FtsZ is derived from the bacterial cell division machinery, whereas cpDRPs belong to the dynamin-related protein family that mediates eukaryotic membrane fission. Together, these distinct GTP-dependent systems operating at multiple sites within the chloroplast division apparatus confer a specialized and complex division mechanism compared with other membrane fission and cytokinetic processes.

FtsZ assembles into filaments through polymerization in the presence of GTP^58^, and these protofilaments bundle to form the FtsZ ring^21,59^ (Supplementary Fig. 3). Assembly of the FtsZ ring is orchestrated by numerous regulatory factors that determine the assembly site and control FtsZ polymerization–depolymerization dynamics^56,60–62^. Some FtsZ regulators reside in the inner chloroplast envelope, serving both as anchoring components for the FtsZ ring and as transducers that convey division-site positional information to the cytosolic side through interactions with outer-envelope proteins^63,64^. Furthermore, heterologous expression studies suggest that the FtsZ ring can generate a driving force for membrane constriction during chloroplast division^21^, analogous to its role in bacterial cytokinesis. However, although FtsZ plays an important role in establishing the chloroplast division site, the contribution of FtsZ to force generation during the constriction step appears to be limited under the *in vitro* conditions used here. Combined with our results, previous findings suggest that the primary role of FtsZ and FtsZ-associated factors may have gradually shifted during evolution from serving as a constriction actuator to regulating chloroplast division-site positioning and division timing in accordance with the cell cycle and developmental stage.

### Evolution of endosymbiotic organelle division rings

Endosymbiotic organelles such as chloroplasts and mitochondria evolved from free-living bacteria through ancient endosymbiotic events. Although bacterial engulfment by a host cell is considered a key trigger of this process, retention of an endosymbiont within the cytosol alone is insufficient to ensure stable inheritance. Without binary fission of the endosymbiont, one of the daughter cells will inevitably lose it during host cell division. Therefore, acquisition of a dedicated division mechanism that ensures faithful segregation represents a pivotal evolutionary transition from endosymbiont to organelle^65^.

Consistent with this hypothesis, division mechanisms of endosymbiotic organelles exhibit notable structural similarities despite their distinct evolutionary histories^1,12,66^. As with the chloroplast division ring, the mitochondrial division ring is composed of mitochondrial FtsZ^67,68^, a DRP^53,69^, and skeletal filaments known as the MD ring^70^, which is assembled by the glycosyltransferase MDR1^34^. Importantly, conservation of these three components is restricted to lower eukaryotes such as *C. merolae*. In contrast, higher eukaryotes, including fungi, animals, and land plants, have lost mitochondrial FtsZ and the MD ring and instead appear to have transitioned to a fission mechanism based on DRP-based filaments. The dynamic cycles of mitochondrial fission and fusion characteristic of higher eukaryotic cells may have imposed selective pressure to simplify the division apparatus, potentially contributing to the emergence of a more efficient and more readily regulatable system. Such a pattern resembles the evolutionary trajectory observed for the chloroplast division machinery^5^.

A striking example of the evolutionary establishment of division machinery for an endosymbiont has been revealed by studies of the trypanosomatid *Angomonas deanei*, an obligate parasite belonging to the clade Discoba^71^. *A. deanei* harbors a single β-proteobacterial endosymbiont per cell, and both FtsZ and DRP are required for endosymbiont division^72^. Unlike the DRP gene, the gene encoding FtsZ is retained in the endosymbiont genome rather than in the host nuclear genome, implying that the endosymbiont in *A. deanei* represents an evolutionary intermediate between a free-living bacterium and an organelle. Although a PD/MD ring-like structure has not yet been identified in *A. deanei*, the requirement for both FtsZ and DRP even at this pre-organelle stage suggests that the shared molecular framework of endosymbiont division may represent an ancestral and broadly conserved mechanism for executing endosymbiotic organelle division.

Building on our findings, investigating division mechanisms in recently described endosymbiotic organelles, such as the nitrogen-fixing nitroplast of *Braarudosphaera bigelowii*^73^ and the photosynthetic chromatophore (cyanelle) of *Paulinella chromatophora*^74^, will be crucial for elucidating the fundamental principles governing endosymbiotic organelle division and organellogenesis.

## Methods

### Cell cultures and synchronous cultivation

The *Cyanidioschyzon merolae* 10D strain (NIES-3377) was used as the wild type in this study^7^. Wild-type and transformed cells were maintained in 2×Allen’s medium [20 mM (NH_4_)_2_SO_4_, 4 mM KH_2_PO_4_, 2 mM MgSO_4_, 1 mM CaCl_2_, 4/1,000 volume of P4 metal solution (0.7 mM FeCl_3_, 0.18 mM MnCl_2_, 77 μM ZnCl_2_, 20 μM CoCl_2_, 10 μM Na_2_MoO_4_, 3 μM Na_2_EDTA), 0.03% (v/v) H_2_SO_4_ to adjust the pH to 2.5]. The uracil auxotrophic M4 strain was maintained in MA2 medium supplemented with 0.5 mg ml^−1^ uracil and 0.8 mg ml^−1^ 5-fluoroorotic acid monohydrate. All strains were cultured in tissue culture flasks (TPP Techno Plastic Products AG, Switzerland) with shaking at 120 rpm under continuous white light (22 μmol m^−2^ s^−1^) at 42 °C. Synchronization of cultures was performed as follows: Cells in asynchronous cultures in the stationary phase were subcultured to <1 × 10^7^ cells ml^−1^ in 100 ml of medium and subjected to a 12-h light/12-h dark cycle at 42 °C under aeration.

### Production of *C. merolae* transformants

To produce the *C. merolae* cell line expressing Venus-fused Dnm2, a construct containing the *Venus-Dnm2* coding sequence flanked by 1,506 bp of upstream and 1,496 bp of downstream sequences was prepared. A glycine-serine (GS)-linker was inserted between *Venus* codon-optimized for expression in *C. merolae* and *Dnm2*. Using the resulting construct as a template, a DNA fragment for homologous recombination comprising 1,506 bp of the upstream sequence of *Dnm2*, *Venus-Dnm2* in-frame with the sequence encoding the GS linker, 200 bp of the downstream sequence of *TUBB*, the *URA5.3* selection marker, and 1,496 bp of the downstream sequence of *Dnm2* was amplified by PCR. The PCR amplicon was transformed into the chromosomal *Dnm2* (CMN262C/XP_005537380) locus of the uracil auxotrophic mutant strain M4 as described^75–77^. Using the construct as a PCR template, transformants expressing each Venus-fused Dnm2 mutant were generated by site-directed mutagenesis using PCR. Transformants expressing Venus-PDR1 or FtsZ2-Venus were also produced using the same approach. Constructs containing the *Venus-PDR1* coding sequence flanked by 1,831 bp of its upstream and 1,482 bp of its downstream sequences or *FtsZ2-Venus* coding sequence flanked by 500 bp of its upstream and 500 bp of its downstream sequences were transformed into the chromosomal *PDR1* (CMR358C/XP_005538591) or *FtsZ2* (CMO089C/XP_005537502) locus of the uracil auxotrophic mutant strain M4.

To produce a transformed cell line harboring an inducible *Venus*-*Dnm2 chimera*, a nitrogen-source-dependent inducible promoter sequence derived from 800 bp of the upstream sequence of a nitrite reductase gene (*NIR*, CMG021C/XP_005535840)^78^ and the DNA sequence encoding the GTPase domain of Dnm1 (CME019C/XP_005535543, residues 25–328) were utilized. DNA fragments encoding the *NIR* promoter-*Venus*-*Dnm2 chimera* and *URA5.3* genes were integrated into upstream of the chromosomal *ura5.3* of the uracil auxotrophic mutant strain M4. Thus, the original sequence of the chromosomal *Dnm2* locus was maintained in the transformant.

All PCR steps were performed using Platinum SuperFi II DNA Polymerase (Thermo Fisher Scientific). Purification and assembly of DNA fragments were performed using a Wizard SV Gel and PCR Clean-up System (Promega) and a NEBuilder HiFi DNA assembly cloning kit (New England Biolabs). Plasmids used to generate *C. merolae* transformants in this study are listed in Supplementary Table 3.

### Isolation of plastids and division rings

Plastids and division rings were isolated as previously described, with minor modifications^13,31,32^. Briefly, synchronized cells in the dividing phase were harvested by centrifugation at 600*g* for 8 min at room temperature (Avanti JXN-30, Beckman Coulter) using a fixed-angle rotor (JLA-9.1000, Beckman Coulter), resuspended in hypotonic isolation buffer [180 mM sucrose, 20 mM HEPES pH 7.6, 5 mM MgCl_2_, 5 mM KCl, 5 mM EGTA, 1× cOmplete EDTA-free protease inhibitor mixture (Roche), 1 mM DTT], and incubated for 30 min at room temperature. After the centrifugation step was repeated and the cells were resuspended in hypotonic isolation buffer, they were incubated for 10 min at 4 °C. The cells were lysed in a French press G-M (Glen Mills Inc.) at 1750 psi. The lysate was incubated for 60 min on ice with 100 μg ml^−1^ DNase I and layered onto three-step Percoll gradients [80%, 60%, and 40% (v/v) Percoll] dissolved in isotonic isolation medium containing 300 mM sucrose. Following centrifugation at 95,500*g* for 50 min at 4 °C in a swinging-bucket rotor (JS-24.38, Beckman Coulter), intact plastids were collected from the band at the 60–80% Percoll interface. The fraction was washed once with isotonic isolation buffer, centrifuged at 600*g* for 15 min at 4°C, and resuspended in isotonic isolation medium. To observe membrane-free division rings, isolated plastids were collected by centrifugation at 570*g* for 5 min at 4 °C and lysed in sucrose-free isolation buffer containing 0.2% (v/v) Nonidet P-40. After 20 min of incubation, the lysate was layered onto one-step Percoll gradients [40% (v/v) Percoll dissolved in sucrose-free isolation medium] and centrifuged at 13,000*g* for 45 min at 4 °C in a swinging-bucket rotor (JS-24.38, Beckman Coulter). Intact division rings with plastid outer membranes were collected from the band at the lysate-40% Percoll interface. The fraction was washed once with sucrose-free isolation buffer, followed by centrifugation at 7,500*g* for 10 min at 4 °C. To dissolve the plastid outer membranes, the fraction was lysed in sucrose-free isolation buffer containing 100 mM *n*-octyl-β-D-glucopyranoside. The membrane-free division ring remained intact in the buffer for 20 min, after which it began to disintegrate.

### Imaging

Microscopy images were acquired using a home-built system constructed around an Olympus IX83 inverted microscope equipped with a 1.45 NA, 100× oil-immersion objective (UPLAPO 100×OHR; Olympus). The microscope was mounted on a vibration-isolated optical table (AY-1510K4; Meiritz Seiki) and controlled using MetaMorph software (Molecular Devices). Illumination was provided by a mercury lamp (U-HGLGPS; Olympus), a 514-nm solid-state laser (Coherent) for fluorescence polarization microscopy, and a 561-nm solid-state laser (Coherent) for TIRF and *d*STORM imaging. Samples were observed through excitation filters (490–500HQ [Olympus] for Venus and FF01-405/10-25 [Semrock] for chloroplasts), dichroic mirrors (Di03-R514-t1-25×36 [Semrock] for Venus, Di03-R561-t1-25×36 [Semrock] for Alexa Fluor 561, and T455lp [Chroma] for chloroplasts), and emission filters (FF02-531/22-25 [Semrock] for Venus, FF02-585/29-25 [Semrock] for Alexa Fluor 561, and FF02-617/73-25 [Semrock] for chloroplasts). Images were recorded with an ORCA-Fusion BT sCMOS camera (Hamamatsu). The effective pixel size was 65.2 × 65.2 nm.

To acquire time-lapse images, fluorescence images were taken in 2 × 2 binning mode and the sample temperature was maintained at 42 °C using an STXG-IX3WX stage-top incubator (Tokai Hit); the stage drift was corrected using a *Z* drift compensator IX3-ZDC2 module (Olympus). The objective was heated to 40 °C with an objective heater (Tokai Hit). Glass-bottom dishes (IWAKI) and medium were warmed at 40 °C before use.

For TIRF and *d*STORM super-resolution imaging, 5 μl of sample was placed onto a glass-bottom dish and gently covered with a clean coverslip. *d*STORM imaging was performed in STORM buffer containing 10 mM β-mercaptoethylamine, and 2,000–4,000 frames were sequentially recorded with an exposure time of 10 ms.

Photobleaching experiments were performed using a Pixel Illuminator-C (Pinpoint Photonics). Purified Venus proteins were used for calibration. Based on these settings, a region of interest was photobleached using a 514-nm laser at 30% power.

For fluorescence polarization microscopy, the linearly polarized illumination from the 514-nm laser was converted into circular polarization using a zero-order achromatic quarter-wave plate (AQWP05M-580; Thorlabs). The emitted fluorescence was then split by W-VIEW GEMINI image-splitting optics (Hamamatsu) equipped with a polarizing plate beamsplitter for 532 nm (PBSW-532R; Thorlabs).

For imaging, 4 ml of cell culture was added to a glass-bottom dish, which was then left to stand in an incubator for 10 min at 40 °C to allow the cells or isolated plastids to freely fall onto the glass. For time-lapse imaging with GTP or its analog, after 1 ml of medium was removed from the glass-bottom dish on the microscope stage, 1 ml of 10 mM nucleotide solution was gently added, and the sample was mixed by pipetting. Excitation light from the mercury lump was attenuated to ∼25% for single-timepoint imaging and ∼3% for time-lapse imaging.

Immunofluorescence and immunoelectron microscopy were performed as previously described^13,14,34^.

### Image processing and analysis

Images were analyzed using Fiji^79^. Background signals were subtracted from *Z*-stack images using a rolling-ball algorithm (radius = 50). Subsequently, maximum-intensity *Z*-projections were generated. Chlorophyll autofluorescence was removed from each *Z*-projected image by subtracting a median-filtered *Z*-projection (radius = 25).

For *d*STORM super-resolution imaging, raw data files were processed using the ThunderSTORM plugin^80^ in Fiji. The photoelectrons per A/D count and base level were set to the manufacturer-specified values of 0.24 electrons per A/D count and 100 A/D count, respectively. Localizations used for downstream analysis were filtered based on the main peak of the sigma distribution, corrected for drift using cross-correlation, and merged when overlapping.

### Protein expression and purification

A construct expressing His₆-tagged GD-2H or GD-3H was generated using the pQE80L vector (Invitrogen). Proteins were produced in the Escherichia coli host strain XL1-Blue (Agilent), and expression was induced for 2 h by adding 0.1 mM isopropyl-β-D-thiogalactopyranoside. The cells were collected by centrifugation, washed with PBS, and resuspended in lysis buffer [20 mM Tris-HCl (pH 7.5), 150 mM KCl, 1× cOmplete EDTA-free protease inhibitor mixture (Roche), 15 mM imidazole, and 1 mM DTT]. Cells were lysed using a French Press G-M (Glen Mills Inc.) at 10,000 psi and subsequently subjected to brief sonication. Following centrifugation at 13,000*g* for 10 min at 4 °C in an angled rotor (MDX-310, Tomy) to remove debris, the supernatant was filtered through a 0.45-μm PVDF membrane (Millex-HV, Merck). His₆-tagged fusion proteins were purified using a 1-ml HiTrap HP column (Cytiva) on an NGC chromatography system (Bio-Rad). Briefly, the lysate was loaded onto a 1-ml HiTrap HP column equilibrated in TK buffer [20 mM Tris-HCl (pH 7.5) and 150 mM KCl]. The column was washed with TK buffer containing 15 mM imidazole and eluted with TK buffer containing 10–500 mM imidazole. The eluted protein samples were dialyzed overnight against TK buffer using a 20-kDa-MWCO Slide-A-Lyzer™ G2 Dialysis Cassette (Thermo Fisher). The samples were frozen in liquid nitrogen and stored at −80°C until use.

### GTP hydrolysis assay

The GTP hydrolysis rates of 0.5 μM wild-type or mutant GD-3H in reaction buffer [20 mM Tris-HCl (pH 7.5), 150 mM KCl, and 4 mM MgCl₂] containing 1 mM GTP (Fujifilm Wako) and 1 mM DTT were monitored over time at 42°C using a malachite green–based assay kit (MAK113, Sigma-Aldrich). The raw GTPase activity data for each condition is provided in Supplementary Data.

### Analytical gel filtration chromatography

Analytical gel filtration chromatography of purified GD-2H or GD-3H, with or without the corresponding ligand, was performed using an ENrich SEC650 column (Bio-Rad) on an NGC chromatography system (Bio-Rad). For each run, 100 μl of purified GD-2H or GD-3H (WT or mutant; 1 mg/ml) was incubated in reaction buffer [20 mM Tris-HCl (pH 7.5), 150 mM KCl, and 4 mM MgCl₂] with or without the corresponding ligand (0.5 mM) for 1 h at 37°C. After centrifugation at 13,000*g* for 3 min, the supernatant was subjected to gel filtration chromatography. The column was equilibrated with TK buffer before sample loading.

### Parameter sweep and phase-diagram analysis of division-ring constriction

We performed coarse-grained mechanochemical simulations of division-ring constriction using a single collective coordinate *ϕ*(*t*), representing the torsional/coiling degree of freedom. The raw radius was defined as *R*_raw_(*φ*) = *R*_0_ – *C_R_ϕ*. Load evaluation used a clipped radius *R*(*ϕ*) = max (*R*_min_, *R*_raw_). The total torque acting on *ϕ* was

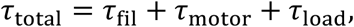

with filament elasticity *τ*_fil_ = −*K*_fil_*ϕ* and motor torque

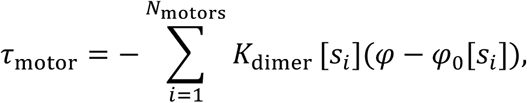

where each motor *i* occupies one of four nucleotide states *s_i_* ∈ {0, 1, 2, 3} (Apo, GTP, GDP·Pi, GDP) with state-dependent preferred angles *ϕ*_0_[*s*]. The load was modeled exclusively as a radial spring of stiffness *K_R_*, implemented via

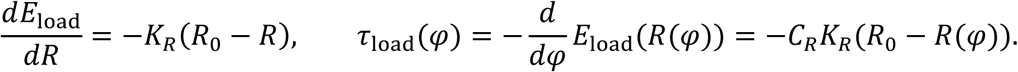

We modeled *ϕ*(*t*) using overdamped Langevin dynamics,

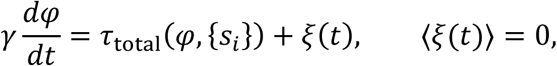

where *τ*_total_ = *τ*_fil_ + *τ*_motor_ + *τ*_load_. For numerical simulations, this stochastic differential equation was integrated with the Euler–Maruyama scheme,

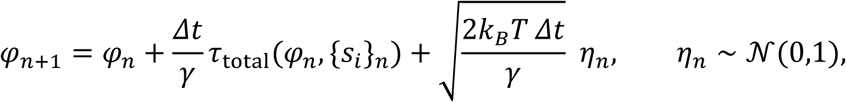

Here, *ξ*(*t*) was taken to be Gaussian white noise with zero mean. In the discrete-time update, this was implemented by drawing an independent standard normal variate *η_n_* at each step (Python: rng.standard_normal()) and scaling it by 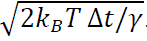. We used Δ*t* = 10^−4^.

Chemical state transitions followed a four-state sequential scheme with rates *k*_on_GTP_, *k*_hyd_, *k*_rel_Pi_, *k*_off_GDP_, applied as stepwise transition probabilities *k*Δ*t*. Scission was defined as the first passage time *t*_sc_ at which *R*_raw_(*ϕ*) < *R*_cut_. The scission success rate and mean scission time were computed over repeated runs (mean time over successful runs only). For phase diagrams, we swept *K_R_* × *γ* and performed 50 independent runs per grid point. To reduce variance in condition comparisons, Condition 1 and Condition 2 were evaluated using identical seed sets at each grid point. The maximum simulation time was set adaptively and smoothly as a function of (*K_R_*, *γ*), with lower and upper caps, to avoid artifacts from piecewise time-window switching.

A detailed description of the coarse-grained mechanochemical model and simulation procedure is provided in the Supplementary Notes.

### Statistical analysis

Statistical comparisons were performed using an unpaired two-sided Student’s *t*-test. The raw data sets and the actual *p* values of Student’s *t*-test are provided in Supplementary Data.

### Data availability

The data supporting the findings of this work are available in the main figures and supplementary information.

### Code availability

Custom code used for simulations and data analysis in this study is available at https://doi.org/10.5281/zenodo.18765746. The full simulation workflow is described in the Supplementary Information, including a detailed model overview and pseudocode.

## Acknowledgements

We thank the members of the Noji PREST (Supra-assembly of biomolecule) for their support and advice during this project.

## Author Contributions

Y.Y. conceived and designed the study, performed all experiments, and analyzed the data. Y.M. generated transformants and measured culture optical density daily to assess growth. Y.Y. and H.K. performed immunoelectron microscopy. Y.M., H.K. and T.K. contributed to the interpretation of the results and helped refine the discussion and conclusions. Y.Y. supervised the project and acquired funding. Y.Y. wrote and revised the manuscript with input from all authors.

## Supplementary Information

**Supplementary Fig. 1.**
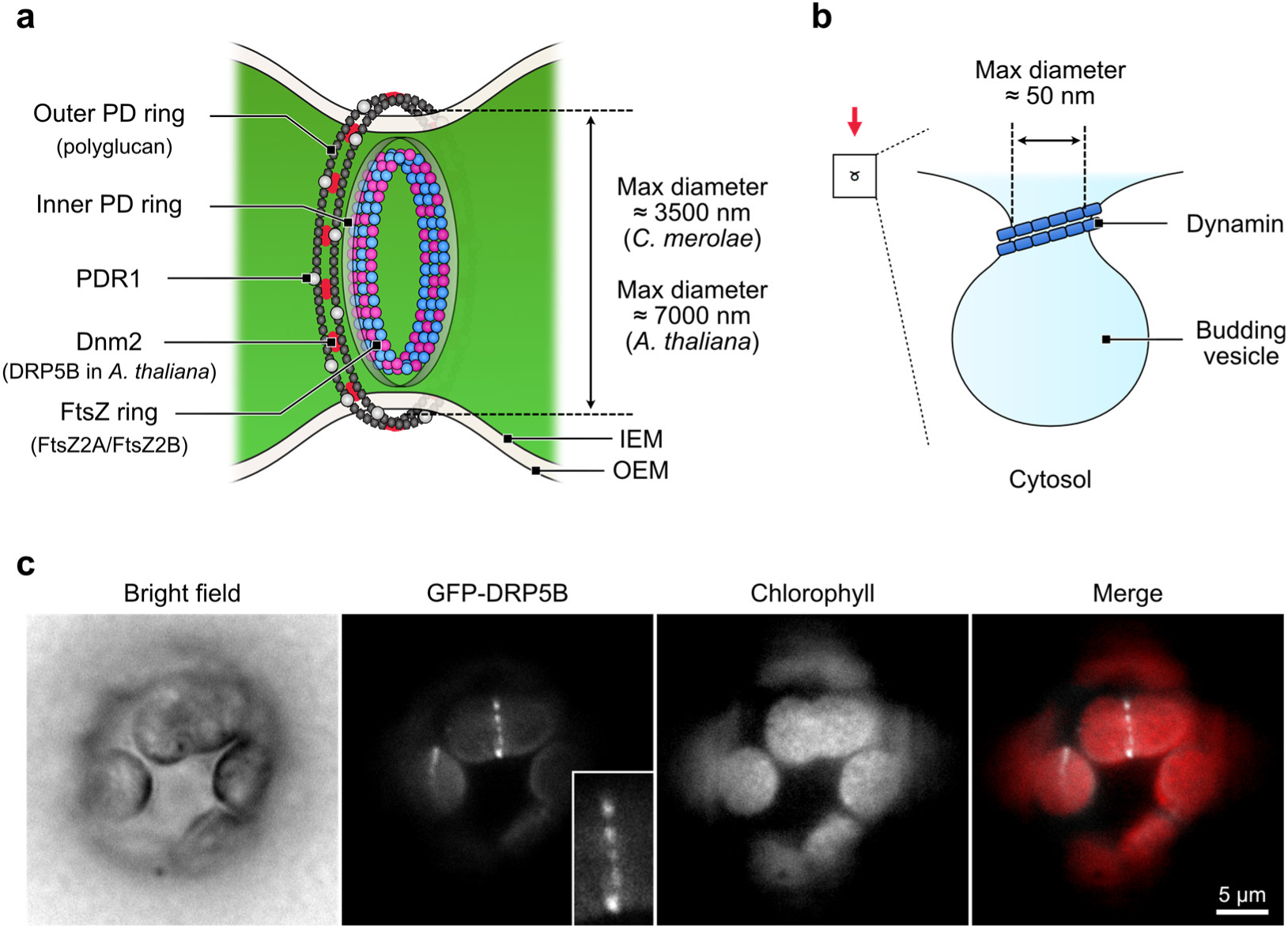
Comparison of chloroplast division machinery and dynamin-based vesicle fission machinery in composition and size scale. **a**, Schematic of the chloroplast division machinery. b, Schematic of the dynamin-based vesicle fission machinery. The vesicle schematic indicated by the red arrow is drawn to the same scale as the dividing chloroplast in (a). c, Fluorescence microscopy images of GFP-tagged DRP5B, a Dnm2 ortholog, in *Arabidopsis thaliana*.

**Supplementary Fig. 2.**
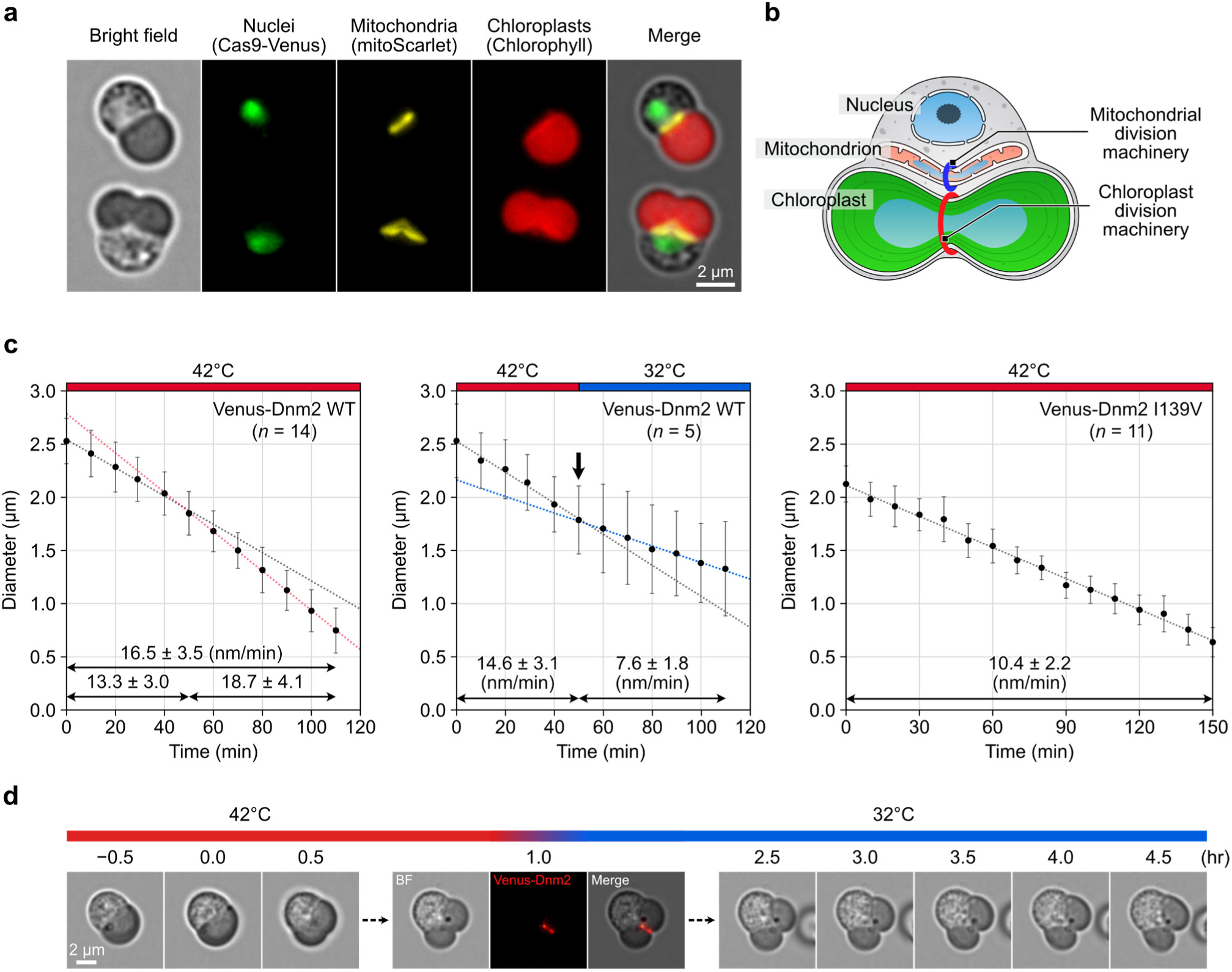
Kinetics of chloroplast division in *C. merolae*. **a**, Representative fluorescence images of *C. merolae* cells. Nuclei (green; Cas9-Venus), mitochondria (yellow; mitoScarlet), and chloroplasts (red; chlorophyll autofluorescence) are shown in a non-dividing cell (top) and a dividing cell (bottom). The *ACTIN* knockout cell was used for the imaging. **b**, Schematic illustration of a dividing *C. merolae* cell. **c**, Time courses of the chloroplast division-plane diameter are shown for Venus-Dnm2 wild-type cells at 42 °C (left), during a temperature shift from 42 °C to 32 °C (middle), and in the Venus-Dnm2 GTPase-domain I139V mutant at 42 °C (right). The timing of the temperature shift is indicated by an arrow. Mean constriction rates are shown in each plot. Constriction rates (nm/min) were calculated as the decrease in division-plane diameter per minute. **d**, Time-lapse imaging of chloroplast division during a temperature shift from 42°C to 32°C.

**Supplementary Fig. 3.**
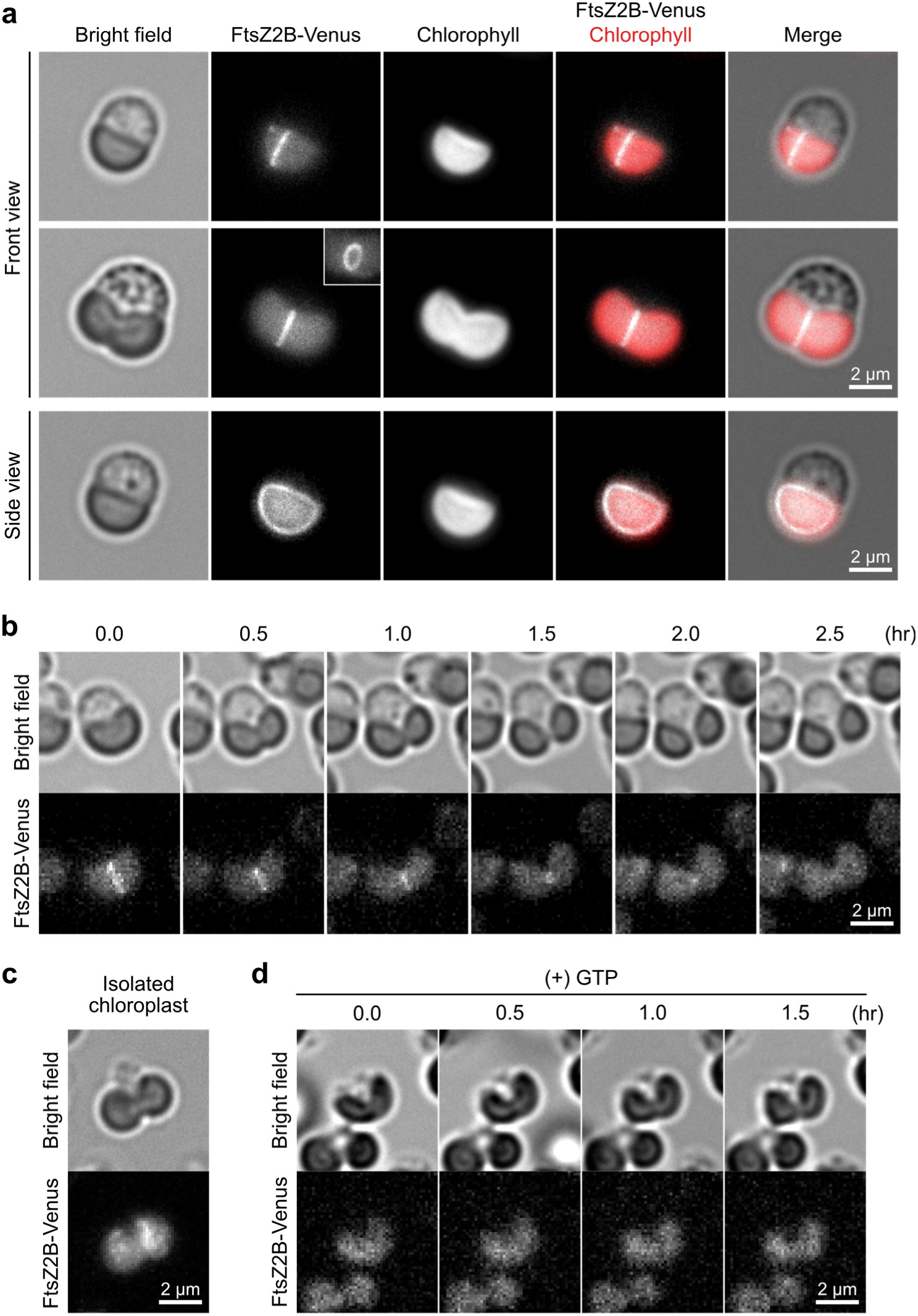
Spatiotemporal dynamics of chloroplast FtsZ during chloroplast division in cells and isolated chloroplasts. **a**, Fluorescence images of cells expressing Venus-tagged FtsZ2B. **b**, Time-lapse imaging of cells expressing FtsZ2B-Venus during chloroplast division. Scale bars, 2 μm. **c**, Representative fluorescence image of an isolated dividing chloroplast expressing FtsZ2B-Venus. **d**, Time-lapse imaging of an isolated chloroplast undergoing division in the presence of 2.5 mM GTP.

**Supplementary Fig. 4.**
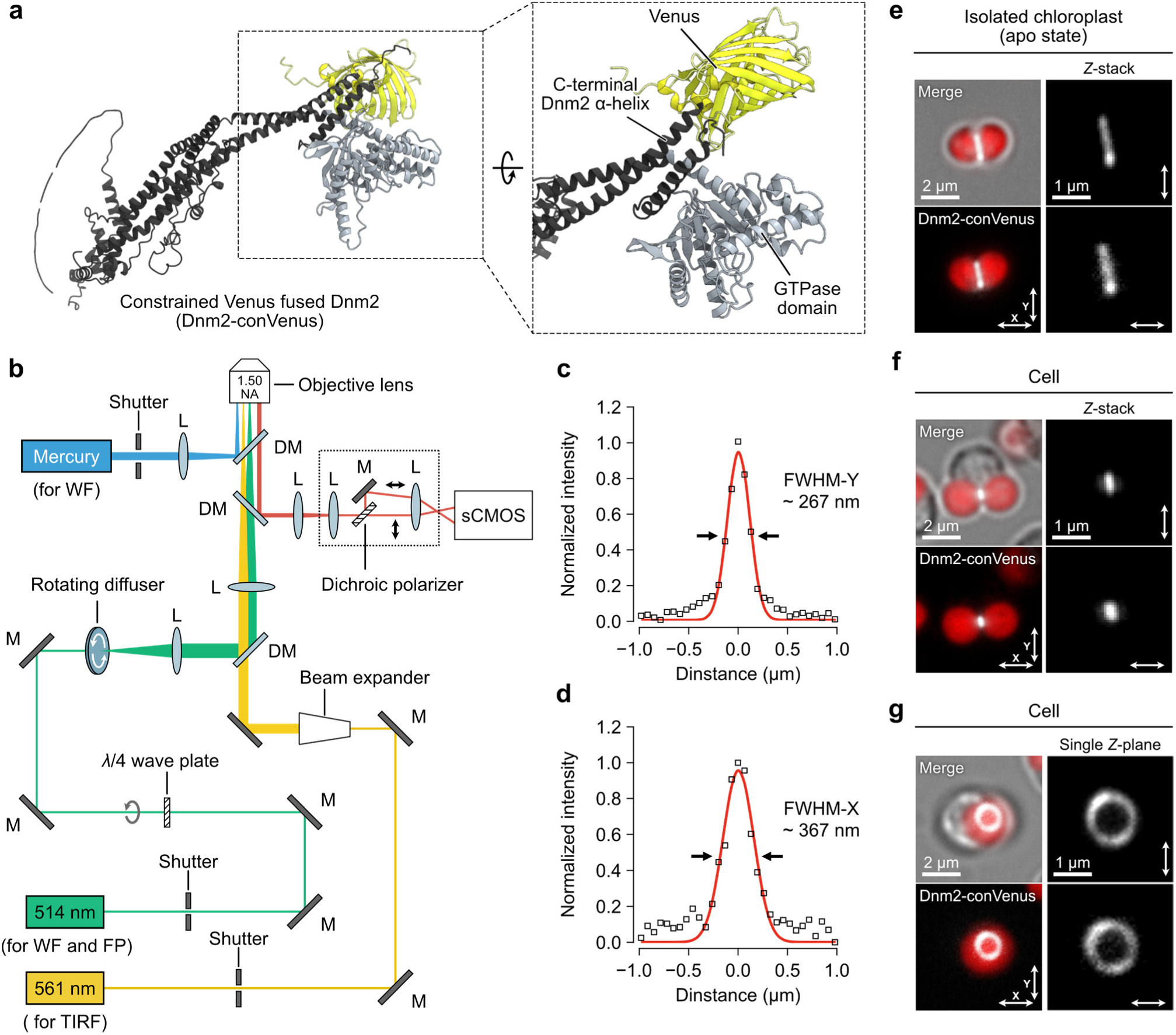
Molecular orientation of Dnm2 on the chloroplast division ring. **a**, Simulated structure of Venus-tagged Dnm2 with a constrained molecular orientation (Dnm2-conVenus). **b**, Schematic of the experimental setup for TIRF, *d*STORM, and fluorescence polarization imaging. Illumination was provided by a 514 nm solid-state laser (green line) for wide-field (WF) and fluorescence polarization (FP) imaging, a 561 nm solid-state laser (yellow line) for TIRF and *d*STORM, and a mercury lamp (blue line) for imaging chlorophyll autofluorescence. Fluorescence from Dnm2-conVenus (red line) was separated by a dichroic polarizer and detected with an sCMOS camera. **c**,**d**, Fluorescence intensity profiles and Gaussian fits obtained by polarized fluorescence microscopy of Dnm2-conVenus. Cross-sectional intensity profiles of the images in Figure 3a along the indicated arrows are shown. Gaussian fits and the corresponding FWHM values are shown for fluorescence polarized along the *Y*-axis (**c**) and the *X*-axis (**d**). **e**– **g**, Polarized fluorescence microscopy of Dnm2-conVenus in an isolated dividing chloroplast (**e**), in cells during the late phase of chloroplast division (**f**), and in cells imaged from a lateral view of the division site (**g**). Images were acquired using parallel polarizers aligned along the *X-* and *Y*-axes (bidirectional arrows).

**Supplementary Fig. 5.**
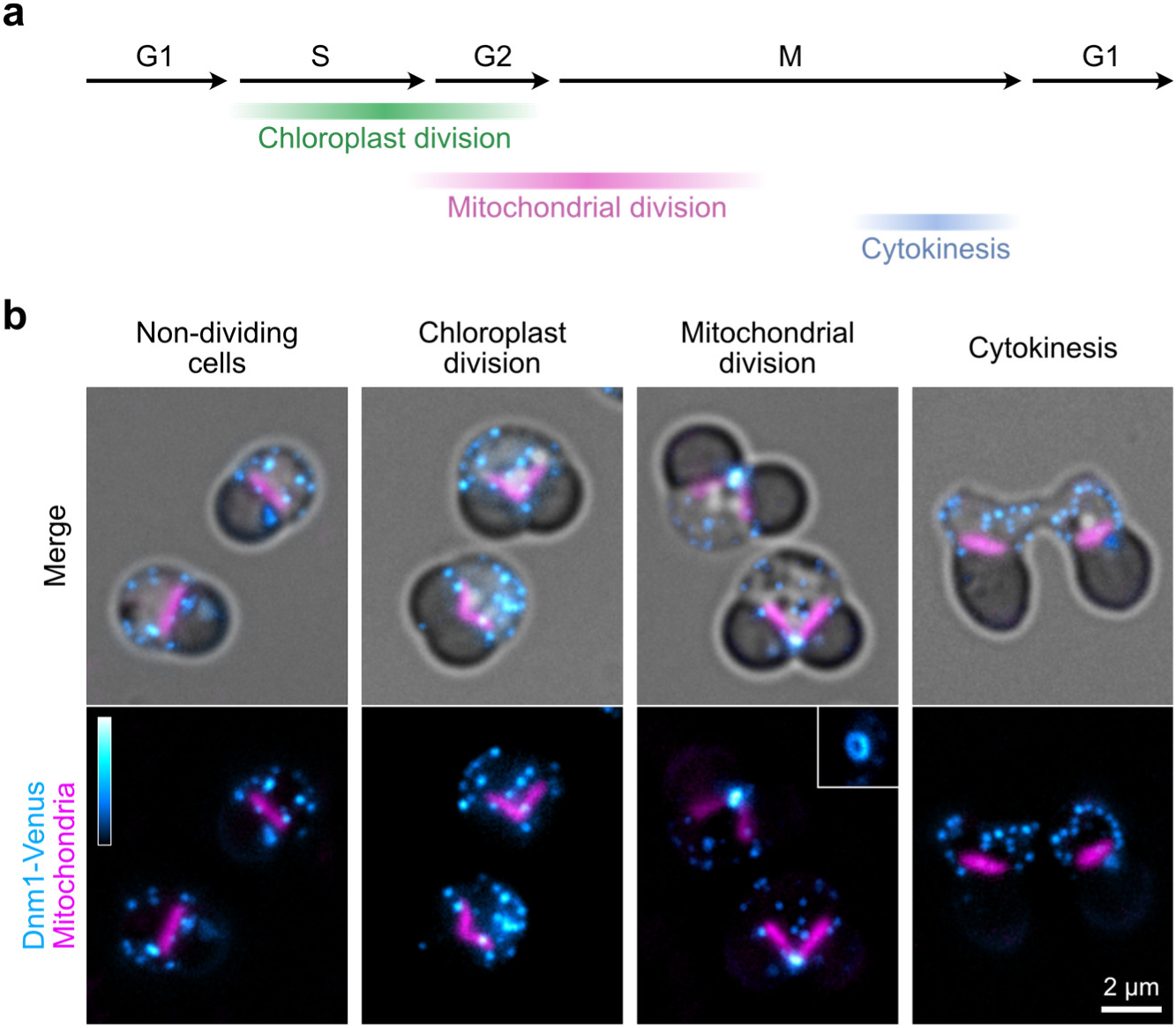
Subcellular localization pattern of Dnm1 in *C. merolae*. **a**, Schematic illustration of the timing of chloroplast division, mitochondrial division, and cytokinesis during the cell cycle. **b**, Dnm1 localization throughout the cell-division cycle. A gene cassette encoding Venus-fused Dnm1 was integrated into the *Dnm1* locus by homologous recombination. Consistent with previous reports^53,81^, we observe that Dnm1-Venus (cyan) appears as >10 cytosolic puncta per cell throughout G1–S phase. From prophase to prometaphase, Dnm1 puncta accumulate and assemble into a ring at the mitochondrial division site (mitochondrion, purple). After mitochondrial division, Dnm1 disperses and returns to a punctate cytosolic distribution. Mitochondria are visualized with MitoTracker Red.

**Supplementary Fig. 6.**
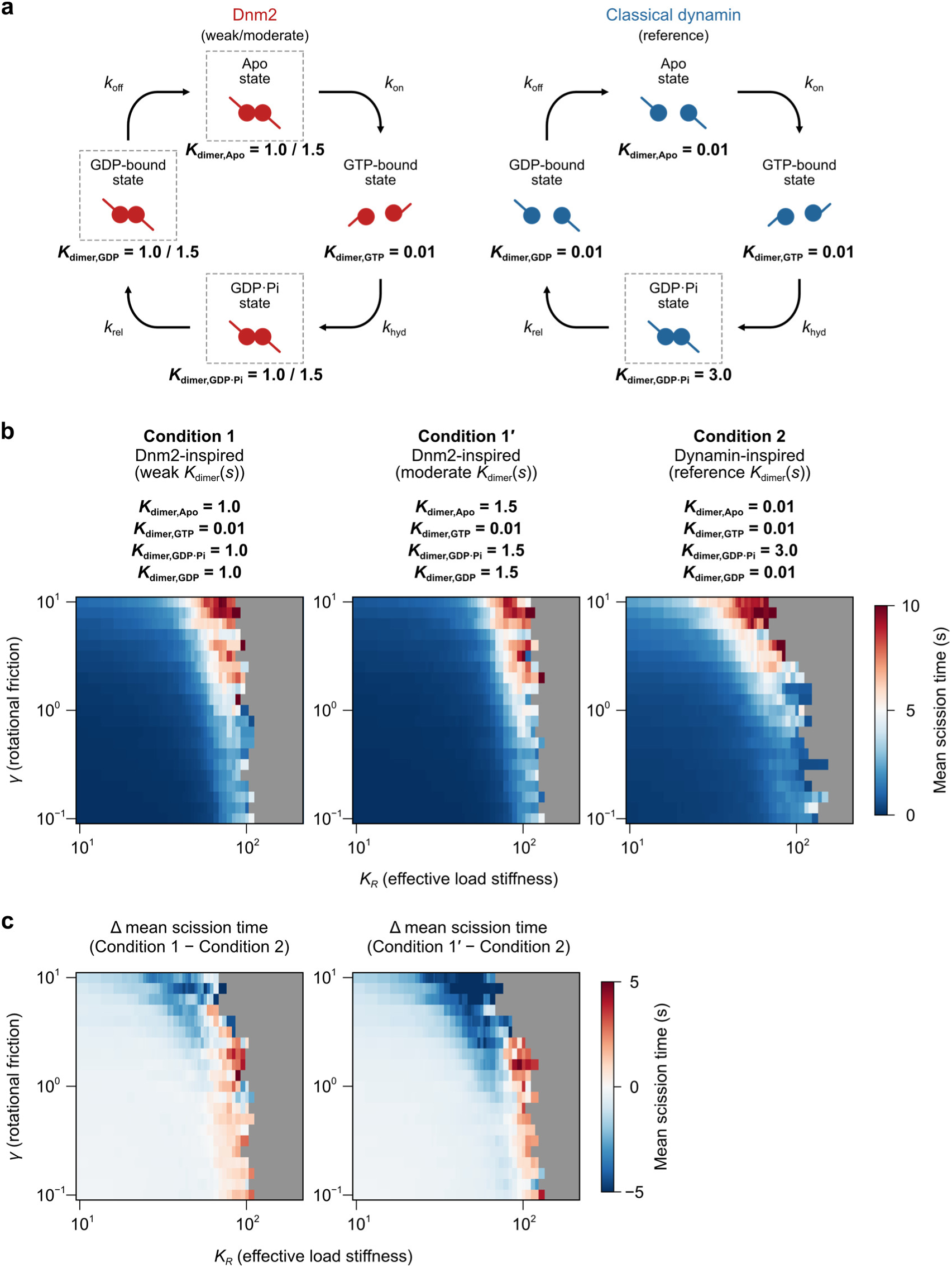
Comparative parameterization of Dnm2-inspired and dynamin-inspired constriction models. **a**, Chemical-state transitions follow a four-state sequential scheme with rates *k*_on_GTP_, *k*_hyd_, *k*_rel_Pi_, *k*_off_GDP_. In the Dnm2-inspired scheme, dimer coupling extends across the GDP·Pi, GDP, and apo states, whereas in the dynamin-inspired scheme strong coupling is largely restricted to the GDP·Pi state. States featuring GTPase–GTPase dimerization are boxed with dotted lines. Based on partial dimerization observed by size-exclusion chromatography (SEC), dimer-coupling stiffness in the Dnm2-inspired scheme was reduced such that *K*_dimer_ in the GDP·Pi/GDP/apo states was set to one-third (Condition 1) or one-half (Condition 1′) of the GDP·Pi-state *K*_dimer_ used in the dynamin-inspired scheme (Condition 2). **b**, Mean time-to-scission phase diagrams for the Dnm2-inspired schemes (Conditions 1 and 1′) and the dynamin-inspired scheme (Condition 2). Regions where scission failed are shown in gray. **c**, Differential mean time-to-scission map (Δ mean scission time) highlighting parameter regimes in which post-hydrolysis coupling improves scission robustness.

## Supplementary Notes

## 1. Model architecture

### 1.1 Model overview

We implemented a coarse-grained mechanochemical model in which division-ring constriction is represented by a single collective coordinate, *ϕ*(*t*), corresponding to the ring’s torsional/coiling degree of freedom. The ring radius is coupled to *ϕ* via a linear geometric relation,

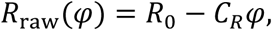

and a clipped radius *R*(*φ*) = max (*R*_min_, *R*_raw_) was used for load calculations. The system comprises *N*_motors_ identical mechanochemical motor units (“motors”; *N*_motors_ = 50), each occupying one of four nucleotide states: Apo (state 0), GTP (state 1), GDP·Pi (state 2), and GDP (state 3). Motors generate state-dependent restorative torques that bias *ϕ* toward a preferred angle *ϕ*_0_[*s*] with an effective stiffness *K*_dimer_[*s*].

### 1.2 Torque decomposition

At any time, the total torque acting on *ϕ* is the sum of (i) a filament torsional elasticity term, (ii) the motor-generated term, and (iii) a load torque originating from a radial spring:

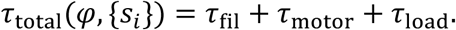

Filament elasticity was modeled as

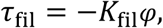

with *K*_fil_ = 0.5.

For the motor term, motor *i* in state *s_i_* ∈ {0, 1, 2, 3} contributes a torque proportional to the deviation from its state-dependent preferred angle *ϕ*_0_[*s*]:

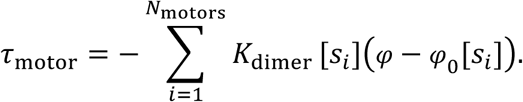

Preferred angles were fixed to *ϕ*_0_ = {2.3, 0.0, 2.9, 2.3} rad for states {0, 1, 2, 3}, respectively. Load was modeled exclusively as a radial spring (radius-spring-only load) with stiffness K_R_, expressed through the radial energy gradient

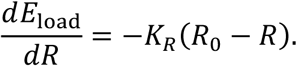

The load torque was computed from a radial spring energy using the chain rule:

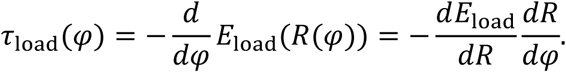

We used the geometric coupling *R*_raw_(*φ*) = *R*_0_ – *C_R_ϕ* and evaluated the load using the clipped radius

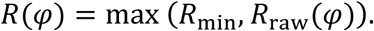

Because *dR*_raw_/*dϕ* = −*C_R_*, the load torque becomes

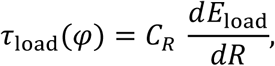

with *dE*_load_/*dR* evaluated at *R* = *R*(*ϕ*). The radial spring was defined by

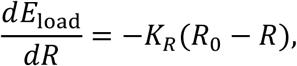

yielding the explicit expression used in simulations:

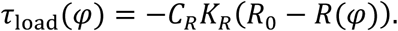

### 1.3 Stochastic dynamics of *ϕ* (overdamped Langevin update)

We modeled *ϕ*(*t*) using overdamped Langevin dynamics,

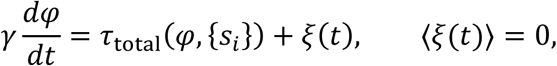

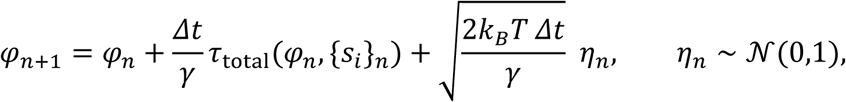

Here, *ξ*(*t*) was taken to be Gaussian white noise with zero mean. In the discrete-time update, this was implemented by drawing an independent standard normal variate *η_n_* at each step (Python: rng.standard_normal()) and scaling it by 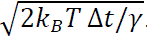. The time step was Δ*t* = 10^−4^, and *k_B_T* = 0.5. Baseline geometric and cutoff parameters were *R*_0_ = 1.0, *C_R_* = 0.32, *R*_cut_ = 0.25, and *R*_min_ = 0.15.

### 1.4 Four-state chemical kinetics

Each motor underwent stochastic transitions among the four nucleotide states in discrete time. At each step, a motor in state *s* transitions to the next state with probability *k*Δ*t*, where the rate constants (s^−1^) were:

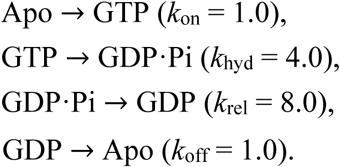

State updates were performed for all motors at every integration step.

### 1.5 Scission criterion and outcome metrics

A trajectory was classified as successful scission when the raw radius satisfied

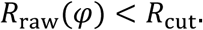

The scission time *t*_sc_ was recorded as the first time the criterion was met. For repeated simulations, the scission success rate was defined as the fraction of runs achieving scission, and the mean scission time was computed over successful runs only (failed runs were excluded).

### 1.6 Simulation conditions (Condition 1 vs Condition 2)

We compared two mechanochemical conditions that differ only in the state dependence of the motor stiffness K_dimer_[s].

**Condition 1 (series)**:

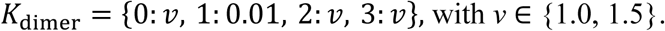

**Condition 2 (fixed)**:

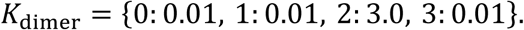

All other parameters were identical between conditions.

### 1.7 Adaptive simulation time window (smooth *T*_max_ rule)

To balance computational cost with accurate estimation of scission outcomes across the (*K_R_*, *γ*) parameter sweep, we used an adaptive maximum simulation time *T*_max_ that increases smoothly with rotational friction *γ* and load stiffness *K_R_*. This prevents slow-but-successful trajectories from being misclassified as failures under a fixed time horizon, while avoiding excessive runtimes in regions where scission is unlikely. Specifically, for each grid point we set

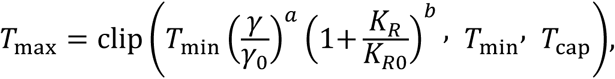

with *T*_min_ = 5.0, *T*_cap_ = 15.0, *K_R_*_0_ = 30.0, *γ*_0_ = 1.0, *a* = 0.7, *b* = 0.5. The number of steps was *N*_steps_ = [*T*_max_/Δ*t*]. We evaluated whether time truncation could bias scission outcomes by checking representative parameter points (including those where *T*_max_ reached *T*_cap_) and found no indication that the main phase-diagram trends were driven by the time-window definition.

### 1.8 Phase-diagram parameter sweeps and variance reduction

Phase diagrams were computed over the *K_R_* × *γ* plane. *γ* was sampled logarithmically from 0.1 to 10 (21 points). *K_R_* was sampled on a nonuniform grid spanning 10 to 200 with denser coverage in intermediate ranges. At each (*K_R_*, *γ*) point, we performed 50 independent runs.

To reduce estimator variance when comparing Condition 1 vs Condition 2, we used common random numbers: the same set of random seeds was used for both conditions at each (*K_R_*, *γ*) point. Specifically, a base seed was assigned per grid point and then incremented to generate a sequence of 50 seeds for that point; these identical seed sequences were used for Condition 1 and Condition 2. We reported condition-specific phase maps for scission success rate and mean scission time, as well as difference maps (Condition 1 – Condition 2).

## 2. Pseudocode describing the mechanochemical simulation and phase-diagram computation

### 2.1 Coarse-grained mechanochemical model

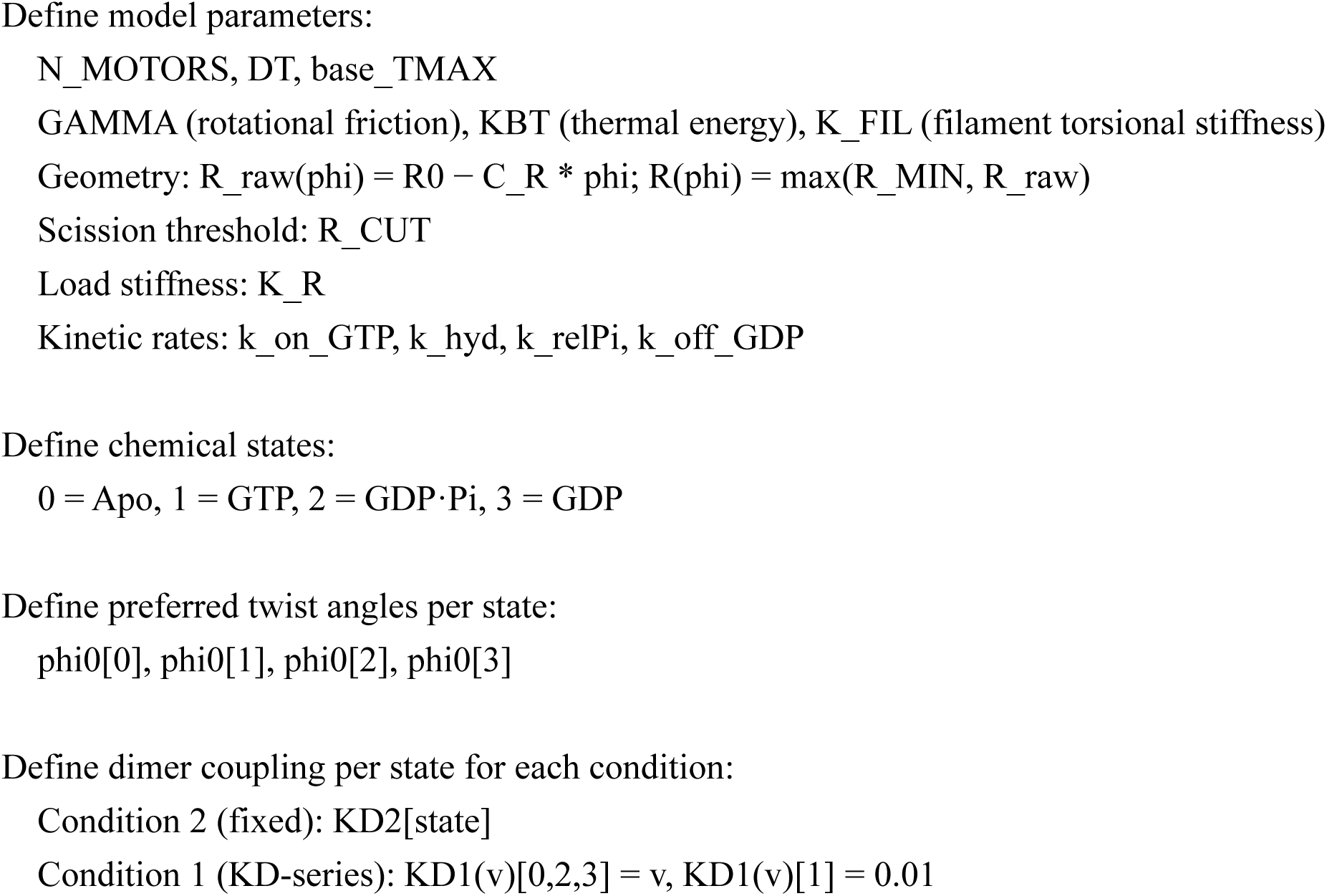

### 2.2 External load model and torque formulation

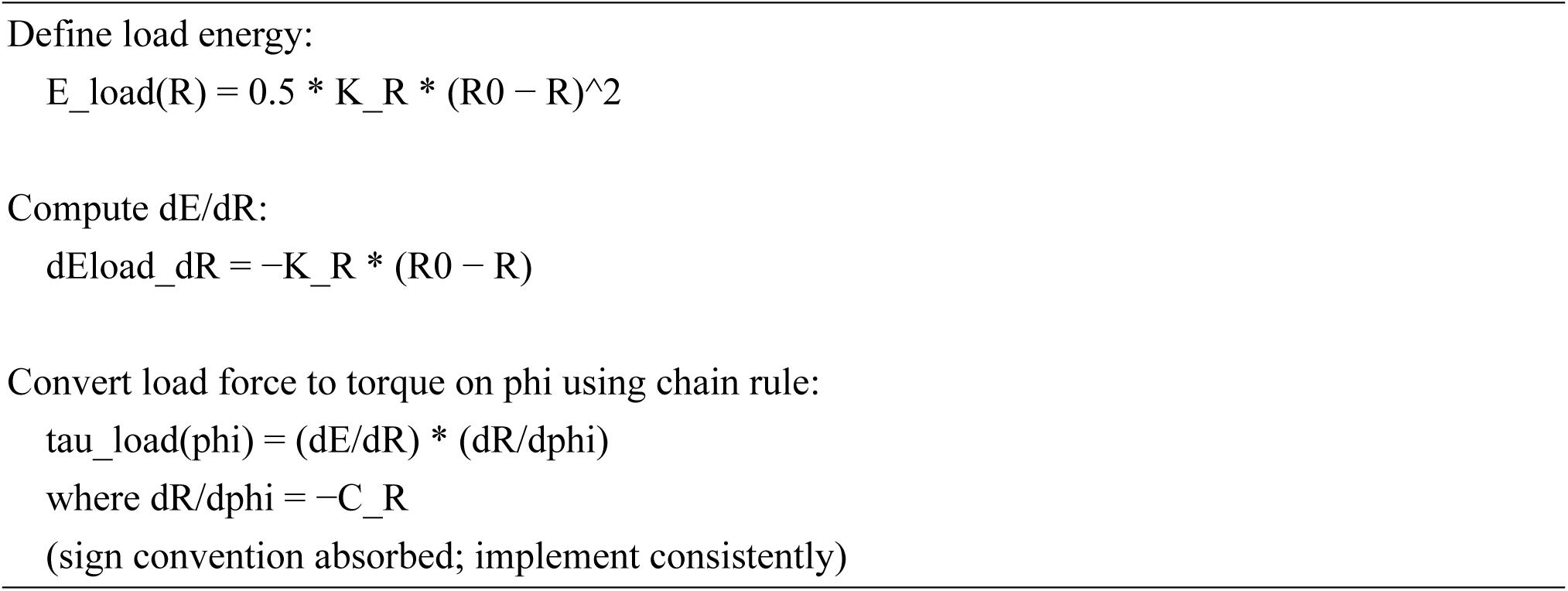

### 2.3 Adaptive time-window scaling

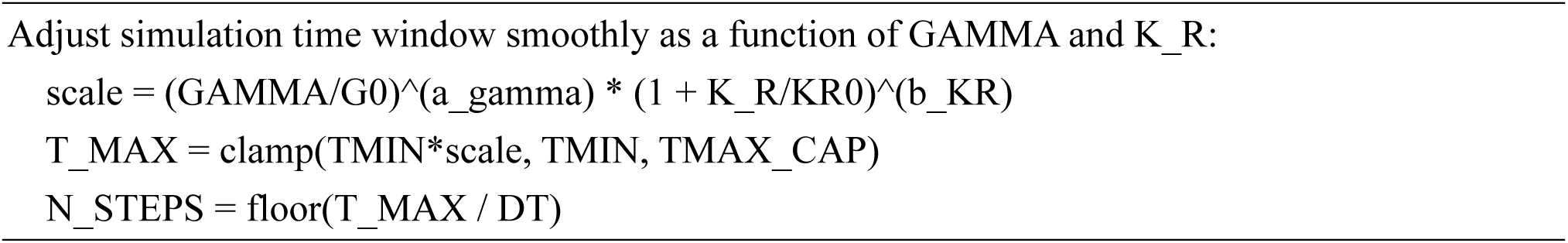

### 2.4 Stochastic chemical state transitions

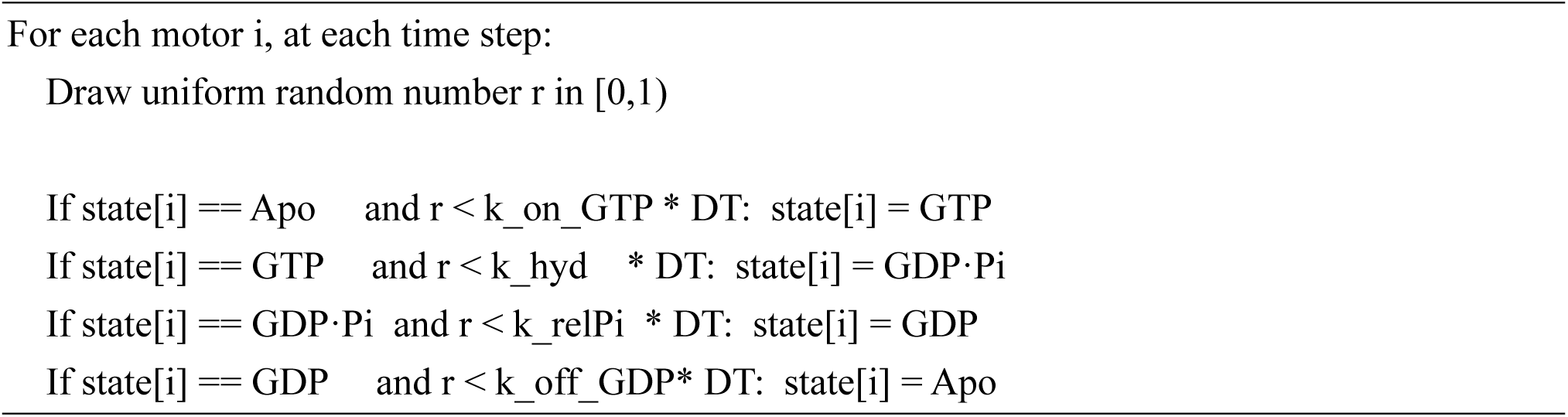

### 2.5 Torque decomposition and force balance

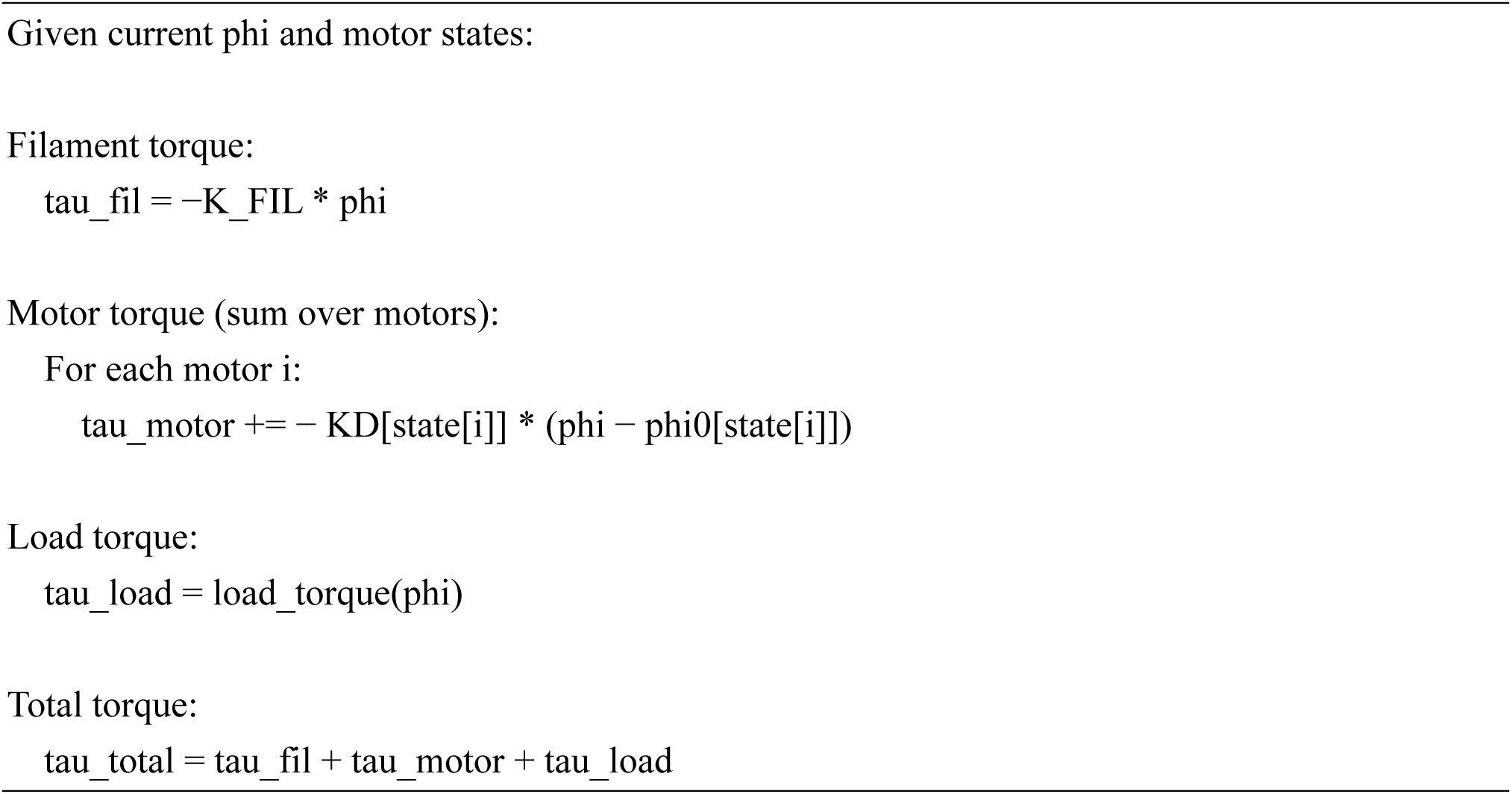

### 2.6 Langevin simulation of ring constriction

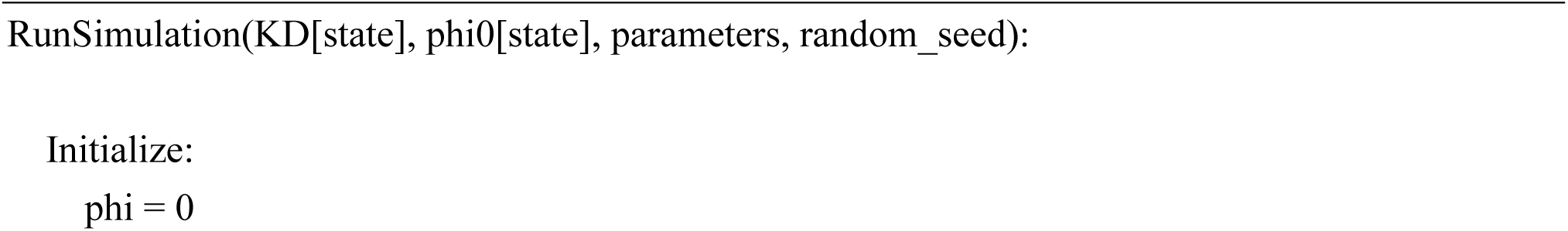
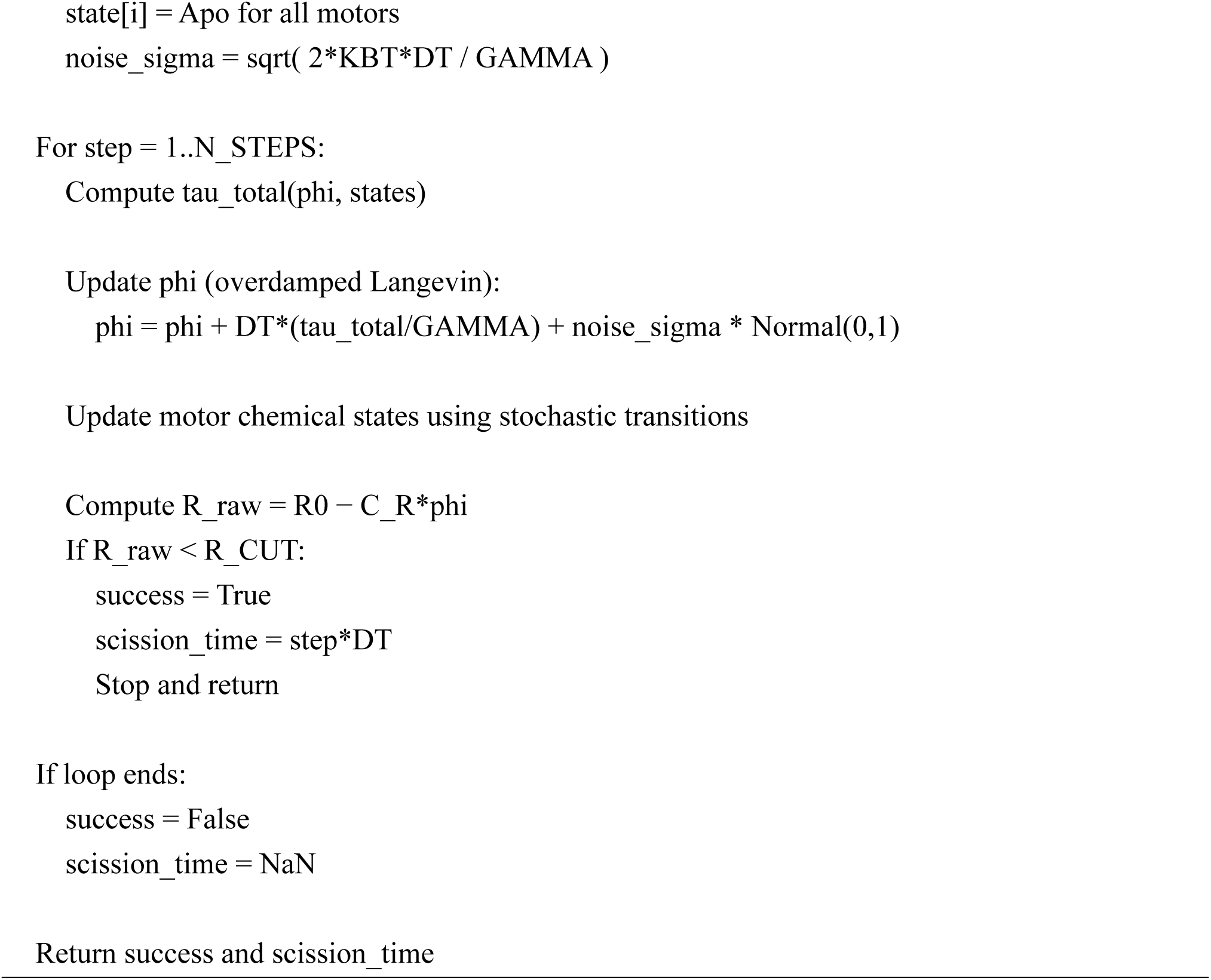

### 2.7 Paired phase-diagram evaluation with common random seeds

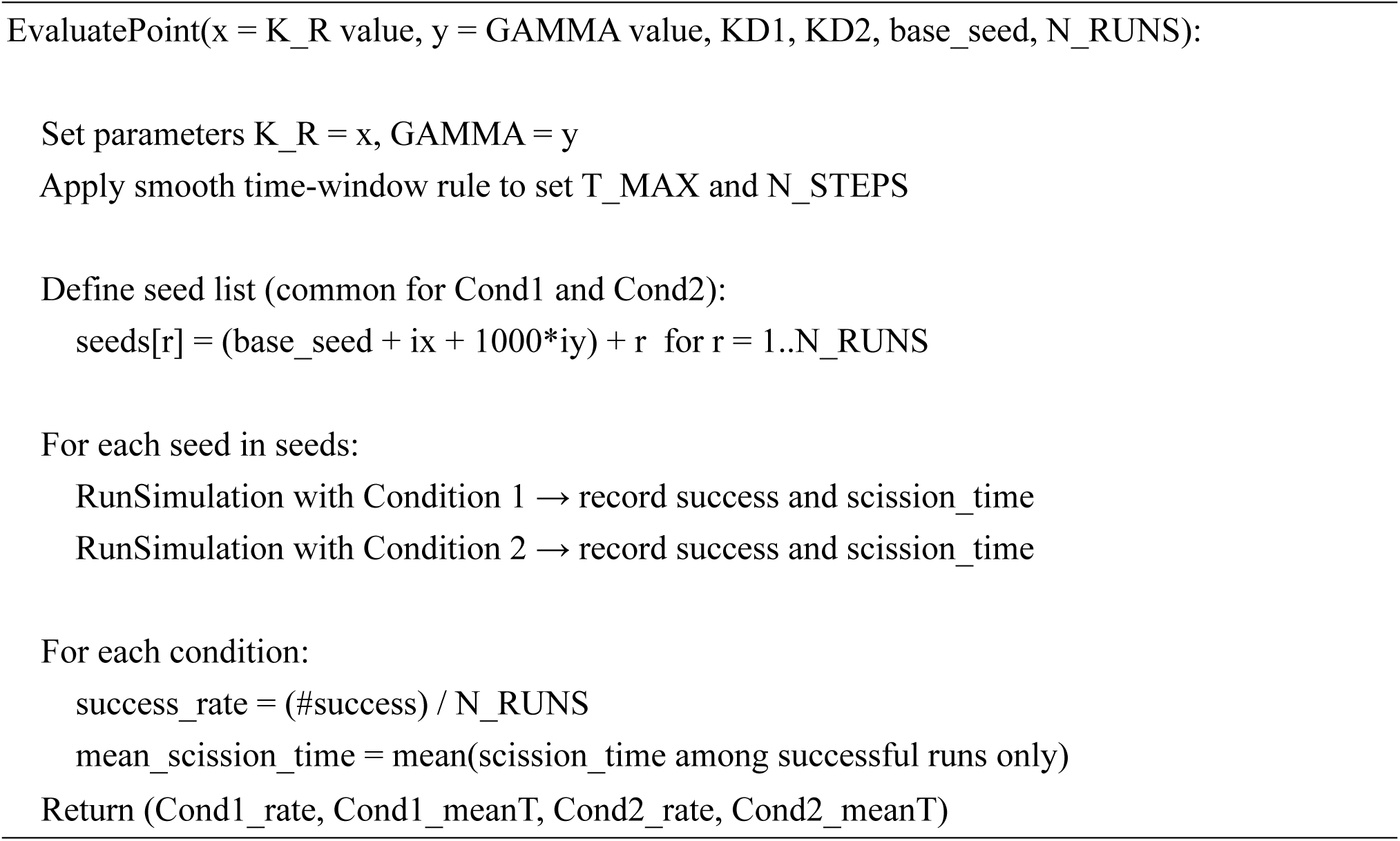

### 2.8 Phase-diagram computation over K_R × GAMMA

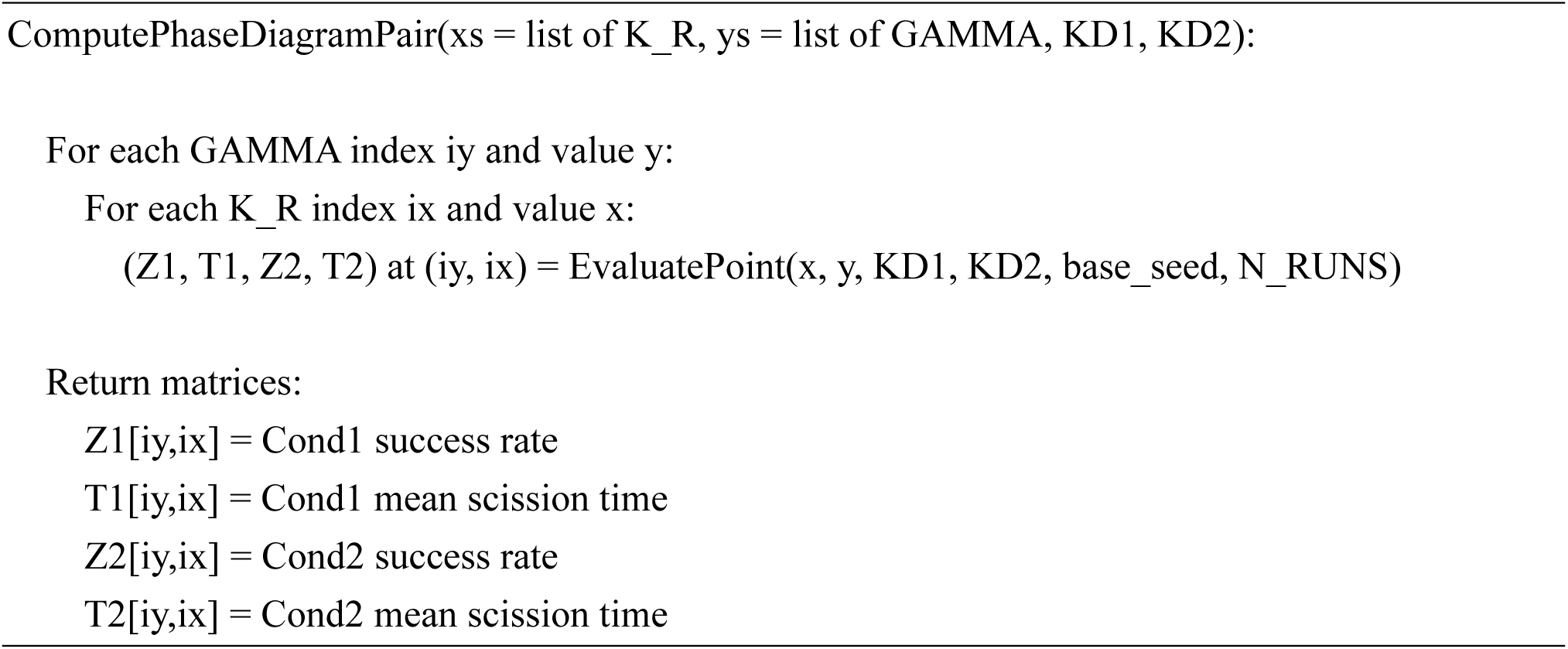

### 2.9 KD-series parameter sweep and differential analysis

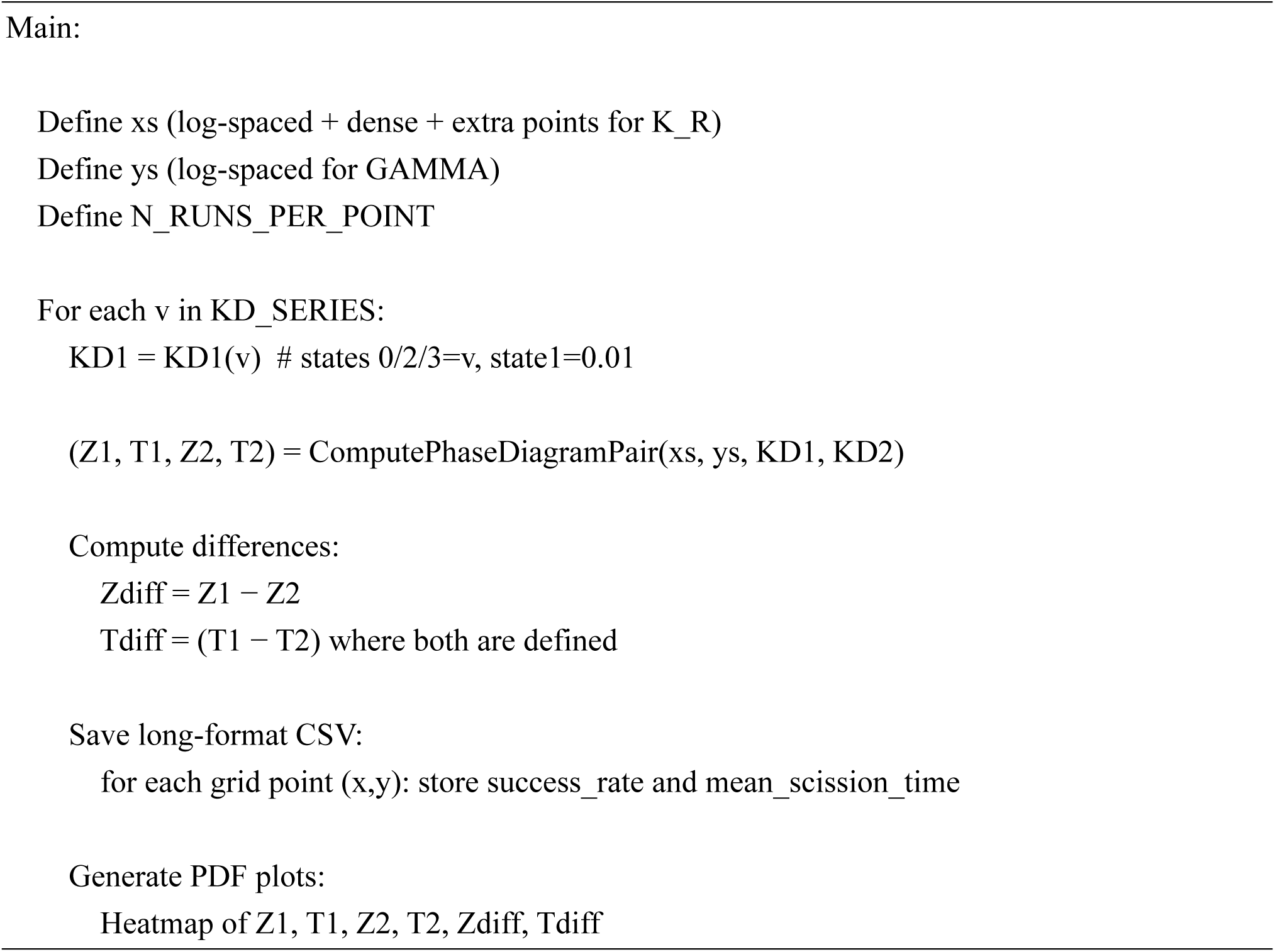

**Supplementary Table 1.**
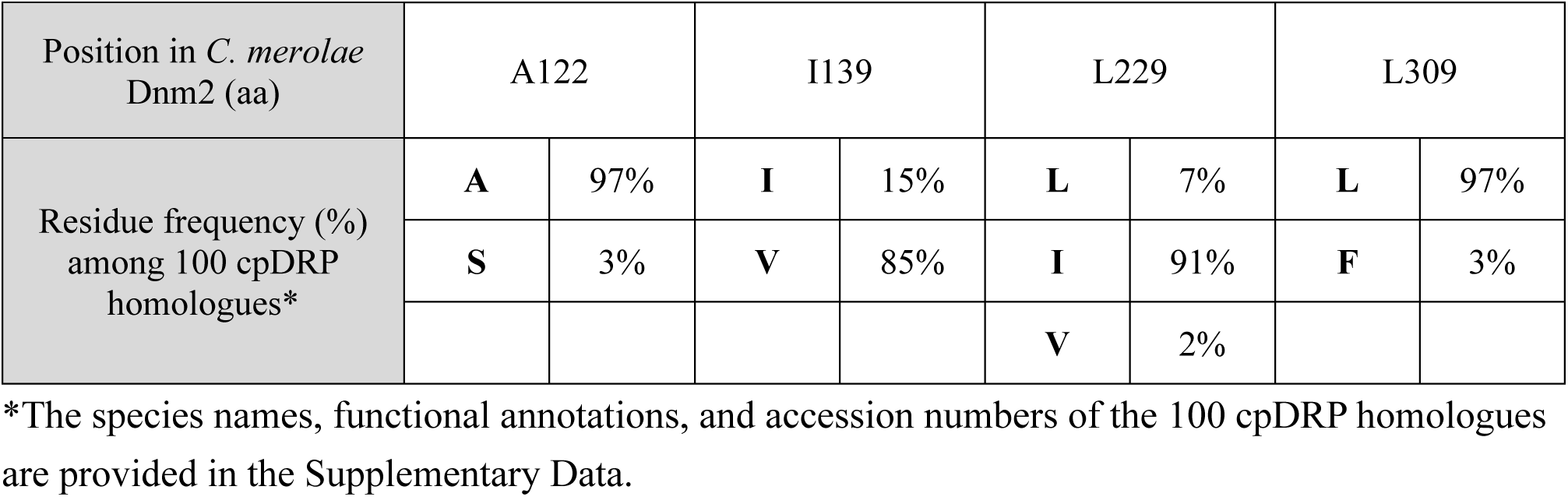
Residue frequency (%) among 100 cpDRP homologues in the G1–G4 motifs.

**Supplementary Table 2.**
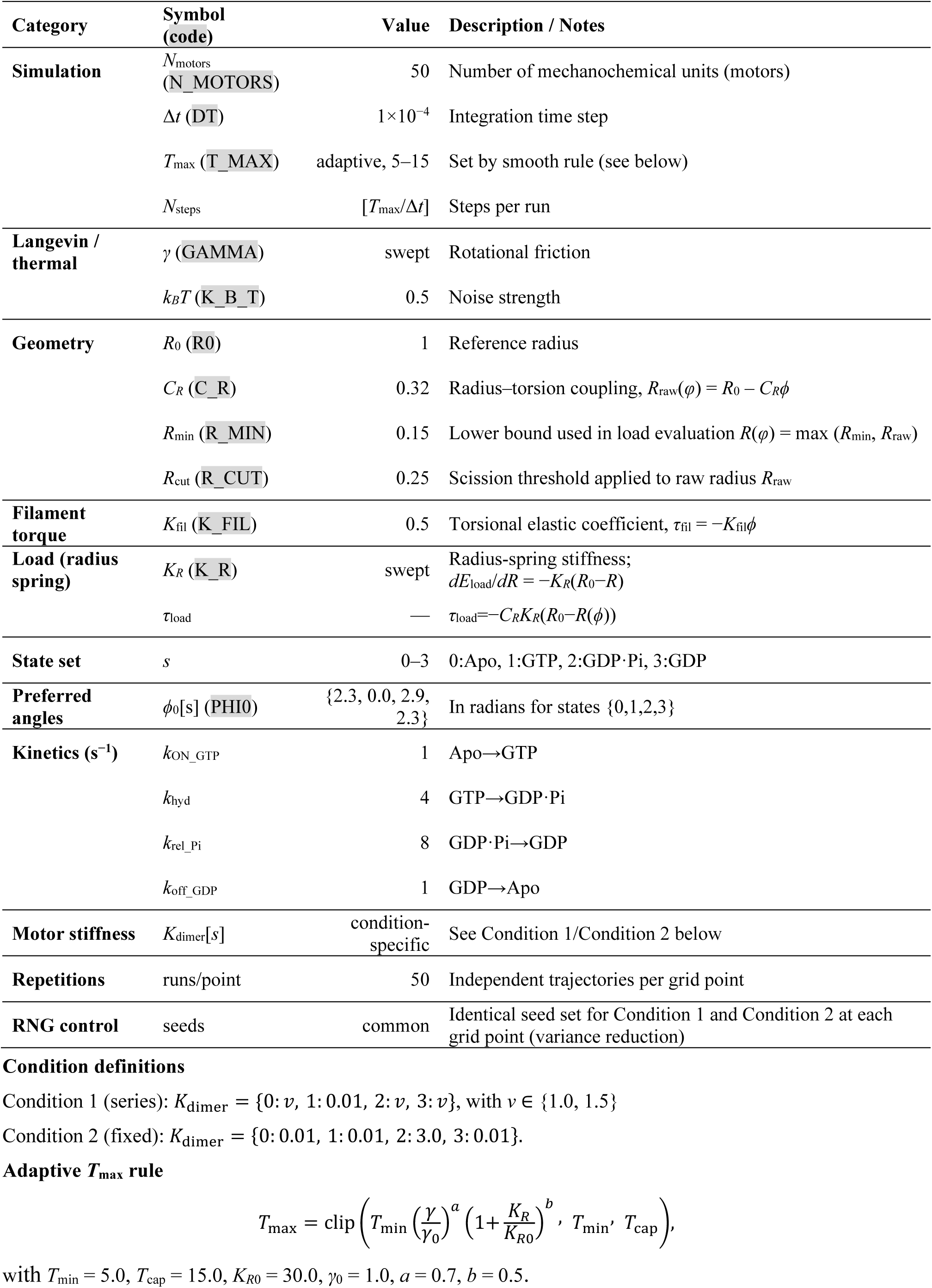
Model parameters and simulation settings.

**Supplementary Table 3.** The plasmids used in this study.

| # | Name | Feature | Source |
| --- | --- | --- | --- |
| 1 | pVenus-Dnm2 | *URA5.3*, *Dnm2* upstream (1506 nt), Venus-Dnm2 | This study |
| 2 | pVenus-Dnm2 A122S | *URA5.3*, *Dnm2* upstream (1506 nt), Venus-Dnm2 A122S | This study |
| 3 | pVenus-Dnm2 I139V | *URA5.3*, *Dnm2* upstream (1506 nt), Venus-Dnm2 I139V | This study |
| 4 | pVenus-Dnm2 L229I | *URA5.3*, *Dnm2* upstream (1506 nt), Venus-Dnm2 L229I | This study |
| 5 | pVenus-Dnm2 L309F | *URA5.3*, *Dnm2* upstream (1506 nt), Venus-Dnm2 L309F | This study |
| 6 | pVenus-Dnm2 chimera | *URA5.3*, *NIR* upstream (800 nt), Venus-Dnm2(1-120)-Dnm1(25-328, K39A)-Dnm2(420-962) | This study |
| 7 | pVenus-PDR1 | *URA5.3*, *PDR1* upstream (1831 nt), Venus-PDR1 | This study |
| 8 | pFtsZ2B-Venus | *URA5.3*, *FtsZ2B* upstream (500 nt), FtsZ2B-Venus | This study |
| 9 | pGD-2H | pQE80L, AmpR, T5 promoter, lac operator, 6xHis, Dnm2(81-445) | This study |
| 10 | pGD-3H | pQE80L, AmpR, T5 promoter, lac operator, 6xHis, Dnm2(81-445)-GSlinker-Dnm2(863-909) | This study |
| 11 | pGD-3H A122S | pQE80L, AmpR, T5 promoter, lac operator, 6xHis, Dnm2(81-445)-GSlinker-Dnm2(863-909) A122S | This study |
| 12 | pGD-3H I139V | pQE80L, AmpR, T5 promoter, lac operator, 6xHis, Dnm2(81-445)-GSlinker-Dnm2(863-909) I139V | This study |
| 13 | pGD-3H L229I | pQE80L, AmpR, T5 promoter, lac operator, 6xHis, Dnm2(81-445)-GSlinker-Dnm2(863-909) L229I | This study |
| 14 | pGD-3H L309F | pQE80L, AmpR, T5 promoter, lac operator, 6xHis, Dnm2(81-445)-GSlinker-Dnm2(863-909) L309F | This study |
| 15 | pGD-3H K135A | pQE80L, AmpR, T5 promoter, lac operator, 6xHis, Dnm2(81-445)-GSlinker-Dnm2(863-909) K135A | This study |
| 16 | pGD-3H chimera | pQE80L, AmpR, T5 promoter, lac operator, 6xHis, Dnm2(81-120)-Dnm1(25-328, K39A)-Dnm2(420-445)-GSlinker-Dnm2(863-909) | This study |

## References

1. Osteryoung, K. W. & Nunnari, J. The division of endosymbiotic organelles. Science 302, 1698–704 (2003).

2. Yoshida, Y., Miyagishima, S.-Y., Kuroiwa, H. & Kuroiwa, T. The plastid-dividing machinery: formation, constriction and fission. Curr. Opin. Plant Biol. 15, 714–721 (2012).

3. Chen, C., MacCready, J. S., Ducat, D. C. & Osteryoung, K. W. The Molecular Machinery of Chloroplast Division. Plant Physiol. 176, 138–151 (2018).

4. Kuroiwa, T. et al. Vesicle, mitochondrial, and plastid division machineries with emphasis on dynamin and electron-dense rings. Int. Rev. Cell Mol. Biol. 271, 97–152 (2008).

5. Miyagishima, S., Nakanishi, H. & Kabeya, Y. Structure, regulation, and evolution of the plastid division machinery. Int. Rev. Cell Mol. Biol. 291, 115–53 (2011).

6. Kuroiwa, T. The primitive red algae Cyanidium caldarium and Cyanidioschyzon merolae as model system for investigating the dividing apparatus of mitochondria and plastids. BioEssays 20, 344–354 (1998).

7. Matsuzaki, M. et al. Genome sequence of the ultrasmall unicellular red alga Cyanidioschyzon merolae 10D. Nature 428, 653–657 (2004).

8. Eberhard, S., Finazzi, G. & Wollman, F. A. The dynamics of photosynthesis. Annu. Rev. Genet. 42, 463–515 (2008).

9. Howe, C. J. J., Barbrook, A. C. C., Nisbet, R. E. R. E. R., Lockhart, P. J. J. & Larkum, A. W. D. W. D. The origin of plastids. Philos. Trans. R. Soc. B Biol. Sci. 363, 2675–2685 (2008).

10. Keeling, P. J. The endosymbiotic origin, diversification and fate of plastids. Philos. Trans. R. Soc. B Biol. Sci. 365, 729–748 (2010).

11. Miyagishima, S. Mechanism of plastid division: from a bacterium to an organelle. Plant Physiol. 155, 1533–44 (2011).

12. Yoshida, Y. & Mogi, Y. How do plastids and mitochondria divide? Microscopy 68, 45–56 (2019).

13. Yoshida, Y. et al. Isolated chloroplast division machinery can actively constrict after stretching. Science 313, 1435–8 (2006).

14. Yoshida, Y. et al. Chloroplasts divide by contraction of a bundle of nanofilaments consisting of polyglucan. Science 329, 949–53 (2010).

15. Bi, E. & Lutkenhaus, J. FtsZ ring structure associated with division in Escherichia coli. Nature 354, 161–164 (1991).

16. Adams, D. W. & Errington, J. Bacterial cell division: assembly, maintenance and disassembly of the Z ring. Nat. Rev. Microbiol. 7, 642–53 (2009).

17. Osteryoung, K. W. & Vierling, E. Conserved cell and organelle division. Nature 376, 473–4 (1995).

18. Mori, T., Kuroiwa, H., Takahara, M., Miyagishima, S. Y. & Kuroiwa, T. Visualization of an FtsZ ring in chloroplasts of Lilium longiflorum leaves. Plant Cell Physiol. 42, 555–9 (2001).

19. Vitha, S., McAndrew, R. S. & Osteryoung, K. W. FtsZ ring formation at the chloroplast division site in plants. J. Cell Biol. 153, 111–20 (2001).

20. Schmitz, A. J., Glynn, J. M., Olson, B. J. S. C., Stokes, K. D. & Osteryoung, K. W. Arabidopsis FtsZ2-1 and FtsZ2-2 are functionally redundant, but FtsZ-based plastid division is not essential for chloroplast partitioning or plant growth and development. Mol. Plant 2, 1211–22 (2009).

21. Yoshida, Y., Mogi, Y., TerBush, A. D. & Osteryoung, K. W. Chloroplast FtsZ assembles into a contractible ring via tubulin-like heteropolymerization. Nat. Plants 2, 16095 (2016).

22. Gao, H., Kadirjan-Kalbach, D., Froehlich, J. E. & Osteryoung, K. W. ARC5, a cytosolic dynamin-like protein from plants, is part of the chloroplast division machinery. Proc. Natl. Acad. Sci. U. S. A. 100, 4328–33 (2003).

23. Miyagishima, S., Froehlich, J. E. & Osteryoung, K. W. PDV1 and PDV2 mediate recruitment of the dynamin-related protein ARC5 to the plastid division site. Plant Cell 18, 2517–30 (2006).

24. Osawa, M., Anderson, D. E. & Erickson, H. P. Reconstitution of contractile FtsZ rings in liposomes. Science 320, 792–4 (2008).

25. Erickson, H. P. Modeling the physics of FtsZ assembly and force generation. Proc. Natl. Acad. Sci. 106, 9238–9243 (2009).

26. Mita, T., Kanbe, T., Tanaka, K. & Kuroiwa, T. A ring structure around the dividing plane of the Cyanidium caldarium chloroplast. Protoplasma 130, 211–213 (1986).

27. Kuroiwa, T. et al. The division apparatus of plastids and mitochondria. Int. Rev. Cytol. 181, 1–41 (1998).

28. Miyagishima, S. Y., Itoh, R., Aita, S., Kuroiwa, H. & Kuroiwa, T. Isolation of dividing chloroplasts with intact plastid-dividing rings from a synchronous culture of the unicellular red alga Cyanidioschyzon merolae. Planta 209, 371–375 (1999).

29. Pyke, K. A. Plastid division and development. Plant Cell 11, 549–556 (1999).

30. Kuroiwa, H., Mori, T., Takahara, M., Miyagishima, S. & Kuroiwa, T. Chloroplast division machinery as revealed by immunofluorescence and electron microscopy. Planta 215, 185–90 (2002).

31. Miyagishima Sy et al. Plastid division is driven by a complex mechanism that involves differential transition of the bacterial and eukaryotic division rings. Plant Cell 13, 2257–68 (2001).

32. Miyagishima, S., Takahara, M. & Kuroiwa, T. Novel filaments 5 nm in diameter constitute the cytosolic ring of the plastid division apparatus. Plant Cell 13, 707–21 (2001).

33. Lomako, J., Lomako, W. M. & Whelan, W. J. Glycogenin: the primer for mammalian and yeast glycogen synthesis. Biochim. Biophys. Acta 1673, 45–55 (2004).

34. Yoshida, Y. et al. Glycosyltransferase MDR1 assembles a dividing ring for mitochondrial proliferation comprising polyglucan nanofilaments. Proc. Natl. Acad. Sci. 114, 13284–13289 (2017).

35. Miyagishima, S. et al. A plant-specific dynamin-related protein forms a ring at the chloroplast division site. Plant Cell Online 15, 655–665 (2003).

36. Praefcke, G. J. K. & McMahon, H. T. The dynamin superfamily: universal membrane tubulation and fission molecules? Nat. Rev. Mol. Cell Biol. 5, 133–47 (2004).

37. Low, H. H. & Löwe, J. A bacterial dynamin-like protein. Nature 444, 766–9 (2006).

38. Ferguson, S. M. & De Camilli, P. Dynamin, a membrane-remodelling GTPase. Nat. Rev. Mol. Cell Biol. 13, 75–88 (2012).

39. Kalia, R. & Frost, A. Open and cut: allosteric motion and membrane fission by dynamin superfamily proteins. Mol. Biol. Cell 30, 2097–2104 (2019).

40. Takei, K., McPherson, P. S., Schmid, S. L. & Camilli, P. De. Tubular membrane invaginations coated by dynamin rings are induced by GTP-γS in nerve terminals. Nature 374, 186–190 (1995).

41. Hopkins, C. R. et al. GTPase activity of dynamin and resulting conformation change are essential for endocytosis. Nature 231–235 (2002) doi:10.1038/35065645.

42. Ford, M. G. J., Jenni, S. & Nunnari, J. The crystal structure of dynamin. Nature 477, 561–6 (2011).

43. Faelber, K. et al. Crystal structure of nucleotide-free dynamin. Nature 477, 556–60 (2011).

44. Chappie, J. S., Acharya, S., Leonard, M., Schmid, S. L. & Dyda, F. G domain dimerization controls dynamin’s assembly-stimulated GTPase activity. Nature 465, 435–440 (2010).

45. Antonny, B. et al. Membrane fission by dynamin: what we know and what we need to know. EMBO J. 35, 2270–2284 (2016).

46. Ford, M. G. J. & Chappie, J. S. The structural biology of the dynamin-related proteins: New insights into a diverse, multitalented family. Traffic 20, 717–740 (2019).

47. Chappie, J. S. et al. A pseudoatomic model of the dynamin polymer identifies a hydrolysis-dependent powerstroke. Cell 147, 209–222 (2011).

48. Ganichkin, O. M. et al. Quantification and demonstration of the collective constriction-by-ratchet mechanism in the dynamin molecular motor. Proc. Natl. Acad. Sci. U. S. A. 118, 1–11 (2021).

49. Yoshida, Y. Insights into the Mechanisms of Chloroplast Division. Int. J. Mol. Sci. 19, 733 (2018).

50. Suzuki, K. et al. Behavior of mitochondria, chloroplasts and their nuclei during the mitotic cycle in the ultramicroalga Cyanidioschyzon merolae. Eur. J. Cell Biol. 63, 280–8 (1994).

51. Fujiwara, T. et al. Periodic gene expression patterns during the highly synchronized cell nucleus and organelle division cycles in the unicellular red alga Cyanidioschyzon merolae. DNA Res. 16, 59–72 (2009).

52. Fujiwara, T., Hirooka, S., Ohbayashi, R., Onuma, R. & Miyagishima, S. Y. Relationship between cell cycle and diel transcriptomic changes in metabolism in a unicellular red alga1[OPEN]. Plant Physiol. 183, 1484–1501 (2020).

53. Nishida, K. et al. Dynamic recruitment of dynamin for final mitochondrial severance in a primitive red alga. Proc. Natl. Acad. Sci. U. S. A. 100, 2146–51 (2003).

54. Chappie, J. S. et al. An Intramolecular Signaling Element that Modulates Dynamin Function In Vitro and In Vivo. Mol. Biol. Cell 20, 3561–3571 (2009).

55. Glynn, J. M. et al. PARC6, a novel chloroplast division factor, influences FtsZ assembly and is required for recruitment of PDV1 during chloroplast division in Arabidopsis. Plant J. 59, 700–11 (2009).

56. Zhang, M., Chen, C., Froehlich, J. E., Terbush, A. D. & Osteryoung, K. W. Roles of Arabidopsis PARC6 in coordination of the chloroplast division complex and negative regulation of FtsZ assembly. Plant Physiol. 170, pp.01460.2015 (2015).

57. Okazaki, K. et al. The PLASTID DIVISION1 and 2 components of the chloroplast division machinery determine the rate of chloroplast division in land plant cell differentiation. Plant Cell 21, 1769–80 (2009).

58. Olson, B. J. S. C., Wang, Q. & Osteryoung, K. W. GTP-dependent heteropolymer formation and bundling of chloroplast FtsZ1 and FtsZ2. J. Biol. Chem. 285, 20634–43 (2010).

59. TerBush, A. D. & Osteryoung, K. W. Distinct functions of chloroplast FtsZ1 and FtsZ2 in Z-ring structure and remodeling. J. Cell Biol. 199, 623–37 (2012).

60. Maple, J., Vojta, L., Soll, J. & Møller, S. G. ARC3 is a stromal Z-ring accessory protein essential for plastid division. EMBO Rep. 8, 293–299 (2007).

61. Maple, J., Aldridge, C. & Møller, S. G. Plastid division is mediated by combinatorial assembly of plastid division proteins. Plant J. 43, 811–23 (2005).

62. Du, W. et al. Enhanced chloroplast FtsZ-ring constriction by the ARC6–ARC3 module in Arabidopsis. Proc. Natl. Acad. Sci. 122, 2017 (2025).

63. Wang, W. et al. Structural insights into the coordination of plastid division by the ARC6-PDV2 complex. Nat. Plants 3, 1–9 (2017).

64. Glynn, J. M., Froehlich, J. E. & Osteryoung, K. W. Arabidopsis ARC6 coordinates the division machineries of the inner and outer chloroplast membranes through interaction with PDV2 in the intermembrane space. Plant Cell 20, 2460–70 (2008).

65. Yoshida, Y. The cellular machineries responsible for the division of endosymbiotic organelles. J. Plant Res. 131, 727–734 (2018).

66. Miyagishima, S., Nishida, K. & Kuroiwa, T. An evolutionary puzzle: chloroplast and mitochondrial division rings. Trends Plant Sci. 8, 432–8 (2003).

67. Takahara, M. et al. A putative mitochondrial ftsZ gene is present in the unicellular primitive red alga Cyanidioschyzon merolae. Mol. Gen. Genet. 264, 452–460 (2000).

68. Beech, P. L. et al. Mitochondrial FtsZ in a Chromophyte Alga. Science (80-.). 287, 1276–1279 (2000).

69. Bleazard, W. et al. The dynamin-related GTPase Dnm1 regulates mitochondrial fission in yeast. Nat. Cell Biol. 1, 298–304 (1999).

70. Kuroiwa, T. et al. Structure, function and evolution of the mitochondrial division apparatus. Biochim. Biophys. Acta 1763, 510–21 (2006).

71. Morales, J. et al. Host-symbiont interactions in Angomonas deanei include the evolution of a host-derived dynamin ring around the endosymbiont division site. Curr. Biol. 33, 28–40.e7 (2023).

72. Maurya, A. K. et al. A nucleus-encoded dynamin-like protein controls endosymbiont division in the trypanosomatid Angomonas deanei. Sci. Adv. 11, 1–17 (2025).

73. Coale, T. H. et al. Nitrogen-fixing organelle in a marine alga. Science (80-.). 384, 217–222 (2024).

74. Nowack, E. C. M. et al. Gene transfers from diverse bacteria compensate for reductive genome evolution in the chromatophore of Paulinella chromatophora. Proc. Natl. Acad. Sci. U. S. A. 113, 12214–12219 (2016).

75. Minoda, A., Sakagami, R., Yagisawa, F., Kuroiwa, T. & Tanaka, K. Improvement of culture conditions and evidence for nuclear transformation by homologous recombination in a red alga, Cyanidioschyzon merolae 10D. Plant Cell Physiol. 45, 667–671 (2004).

76. Ohnuma, M., Yokoyama, T., Inouye, T., Sekine, Y. & Tanaka, K. Polyethylene glycol (PEG)-mediated transient gene expression in a red alga, Cyanidioschyzon merolae 10D. Plant Cell Physiol. 49, 117–20 (2008).

77. Imamura, S. et al. R2R3-type MYB transcription factor, CmMYB1, is a central nitrogen assimilation regulator in Cyanidioschyzon merolae. Proc. Natl. Acad. Sci. 106, 14180–14180 (2009).

78. Fujiwara, T. et al. A nitrogen source-dependent inducible and repressible gene expression system in the red alga Cyanidioschyzon merolae. Front. Plant Sci. 6, 1–10 (2015).

79. Schindelin, J., et al. Fiji: an open-source platform for biological-image analysis. Nat. Methods 9, 676–682 (2012).

80. Ovesný, M., Křížek, P., Borkovec, J., Švindrych, Z. & Hagen, G. M. ThunderSTORM: A comprehensive ImageJ plug-in for PALM and STORM data analysis and super-resolution imaging. Bioinformatics 30, 2389–2390 (2014).

81. Mogi, Y. et al. Genome-wide changes of protein translation levels for cell and organelle proliferation in a simple unicellular alga. Proc. Jpn. Acad. Ser. B. Phys. Biol. Sci. 101, 41–53 (2025).

